# Patterns and determinants of the global herbivorous mycobiome

**DOI:** 10.1101/2022.11.21.517404

**Authors:** Casey H. Meili, Adrienne L. Jones, Alex X. Arreola, Jeffrey Habel, Carrie J. Pratt, Radwa A. Hanafy, Yan Wang, Aymen S. Yassin, Moustafa A. TagElDein, Christina D. Moon, Peter H. Janssen, Mitesh Shrestha, Prajwal Rajbhandari, Magdalena Nagler, Julia M. Vinzelj, Sabine M. Podmirseg, Jason E. Stajich, Arthur L. Goetsch, Jerry Hayes, Diana Young, Katerina Fliegerova, Diego Javier Grilli, Roman Vodička, Giuseppe Moniello, Silvana Mattiello, Mona T. Kashef, Yosra I. Nagy, Joan A. Edwards, Sumit Singh Dagar, Andrew P. Foote, Noha H. Youssef, Mostafa S. Elshahed

**Author notes:** Corresponding author: Address 1110 S. Innovation Way, Stillwater, OK. and.

## Abstract

In spite of their indispensable role in host nutrition, the anaerobic gut fungal (AGF) component of the herbivorous gut microbiome remains poorly characterized. To examine global patterns and determinants of AGF diversity, we generated and analyzed an amplicon dataset from 661 fecal samples from 34 animal species, 9 families, and 6 continents. We identified 56 novel genera, greatly expanding AGF diversity beyond current estimates. Both stochastic (homogenizing dispersal and drift) and deterministic (homogenizing selection) processes played an integral role in shaping AGF communities, with a higher level of stochasticity observed in foregut fermenters. Community structure analysis revealed a distinct pattern of phylosymbiosis, where host-associated (animal species, family, and gut type), rather than ecological (domestication status and biogeography) factors predominantly shaped the community. Hindgut fermenters exhibited stronger and more specific fungal-host associations, compared to broader mostly non-host specific associations in foregut fermenters. Transcriptomics-enabled phylogenomic and molecular clock analyses of 52 strains from 14 genera indicated that most genera with preferences for hindgut hosts evolved earlier (44-58 Mya), while those with preferences for foregut hosts evolved more recently (22-32 Mya). This pattern is in agreement with the sole dependence of herbivores on hindgut fermentation past the Cretaceous-Paleogene (K-Pg) extinction event through the Paleocene and Eocene, and the later rapid evolution of animals employing foregut fermentation strategy during the early Miocene. Only a few AGF genera deviated from this pattern of co-evolutionary phylosymbiosis, by exhibiting preferences suggestive of post-evolutionary environmental filtering. Our results greatly expand the documented scope of AGF diversity and provide an ecologically and evolutionary-grounded model to explain the observed patterns of AGF diversity in extant animal hosts.

## Introduction

Plant biomass represents the most abundant [1], yet least readily digestible [2] nutritional source on Earth. The rise of herbivory in tetrapods was associated with multiple evolutionary innovations to maximize plant biomass degradation efficiency [3, 4]. Extant families of mammalian herbivores are characterized by the enlargement of portions of the hindgut (colon, caecum, or rectum), or the evolution of pregastric structures (diverticula or fermentative sacs in pseudoruminants, and the more complex four-gastric chamber in ruminants) [5, 6]. This allowed for longer food retention times as well as the acquisition and retention of an endosymbiotic anaerobic microbial community, both of which enhance the breakdown of ingested plant material and increase feed energy supply to the host [5, 7]. Within the highly diverse microbial consortia residing in the expanded herbivorous alimentary tract, the anaerobic gut fungi (AGF, phylum Neocallimastigomycota) were the last to be recognized [8-10] and remain the most enigmatic. In spite of their critical role in initiating plant biomass colonization [11, 12], their wide array of highly efficient lignocellulolytic enzymes [13-22], and their biotechnological potential [23-25], AGF diversity and distribution patterns remain, to-date, very poorly characterized [26]. Culture-independent efforts targeting AGF have long been hampered by the documented shortcomings of the universal fungal ITS1 barcoding marker for accurately characterizing AGF diversity [26, 27] and, until recently, by the lack of clear thresholds and procedures for genus and species OTUs delineation [28]. This is reflected in the relatively limited number of high-throughput diversity surveys conducted so far (Table S1). Further, most prior studies were limited in scope and/or breadth, usually analyzing a limited number of samples from a single or few mostly domesticated hosts residing in a single location. Given the large number of extant putative mammalian hosts (e.g. the family Bovidae comprises 8 subfamilies, more than 50 genera, and 143 extant species [29]), as well as the immense number of herbivorous mammals on Earth (a conservative estimate of 75.3 million wild, and 3.5 billion domesticated ruminants, including ≈1.4 billion cattle, 1.1 billion sheep, 0.85 billion goats, ∼60 million horses [30], and ∼50 million donkeys [31]), it is clear that the global AGF diversity remains severely under-sampled.

Beyond documenting diversity and identifying novel lineages, the current patchy and incomplete view of AGF diversity precludes any systematic analysis of the patterns (distribution, relative abundance, and AGF taxa distribution preferences) and determinants (role of and interplay between various factors in structuring communities) of the global herbivorous mycobiome. Assembly and structuring of microbial communities could be governed by deterministic (niche theory-based) or stochastic (null theory-based) processes [32]. The co-occurrence and dynamic interplay between deterministic and stochastic processes is increasingly being recognized [32-34]. Stochastic processes generate changes in community diversity that would not be distinguishable from those changes produced by random chance and include dispersal (movement of organisms from one location to another with subsequent successful colonization in the new location), and drift (defined as random changes in relative abundances of species or individuals due to stochastic factors such as birth, death, or multiplication). Possible deterministic processes governing AGF community assembly include animal host identity (family, species), and gut-type (foregut ruminant, fermenting pseudoruminant, and hindgut fermenters). Beyond host-associated factors, AGF communities could also be impacted by the host domestication status (i.e., whether reared in a domesticated setting and hence predominantly grazers on grasses, or are wild and hence predominantly browsers for fruits, shoots, shrubs, forbs, and tree leaves diets [30]), as well as biogeography, age, sex, or local feed chemistry.

To assess global patterns and determinants of AGF diversity, a consortium of scientists from 16 institutions have sampled fecal material from domesticated and non-domesticated animals from 6 continents covering 9 mammalian families, and 3 gut types. The dataset obtained was used to document the scope of AGF diversity on a global scale and to assess evolutionary and ecological drivers shaping AGF diversity and community structure using the large ribosomal subunit (LSU) as a phylogenetic marker [26]. Furthermore, to assess the evolutionary drivers underpinning the observed pattern of animal host-AGF phylosymbiosis, a parallel transcriptomics sequencing effort for 20 AGF strains from 13 genera was conducted and combined with previous efforts [35-40]. The expanded AGF transcriptomic dataset (52 strains from 14 genera) enabled phylogenomic and molecular timing analysis that correlated observed ecological patterns with fungal and hosts evolutionary histories. Our results greatly expand the scope of documented AGF diversity, demonstrate the complexity of ecological processes shaping AGF communities, and demonstrate that host-specific evolutionary processes (e.g. evolution of host families, genera, and gut architecture) played a key role in driving a parallel process of AGF evolution and diversification.

## Results

### Overview

A total of 661 samples belonging to 34 species and 9 families of foregut-fermenting ruminant (thereafter ruminant, n=468), foregut-fermenting pseudoruminant (thereafter pseudoruminant, n=17), and hindgut fermenters (n=176) were examined (Fig. 1a-b, Table S2). Many of the samples belong to previously unsampled/rarely sampled animal families (e.g. Caviidae, Trichechidae) and species (capybara, mara, manatee, markhor, chamois, takin). The dataset also provides a high level of replication for a variety of animals (229 cattle, 138 horses, 96 goats, 71 sheep, and 23 white-tail deer, among others) (Fig. 1b), locations (418 samples from USA, 74 from Egypt, 38 from Italy, 35 from New Zealand, 31 from Germany, and 25 from Nepal, among others) (Fig. 1a, Table S2), and domestication status (564 domesticated, 97 undomesticated) (Fig. 1b, Table S2), allowing for robust statistical analysis.

**Fig. 1.**
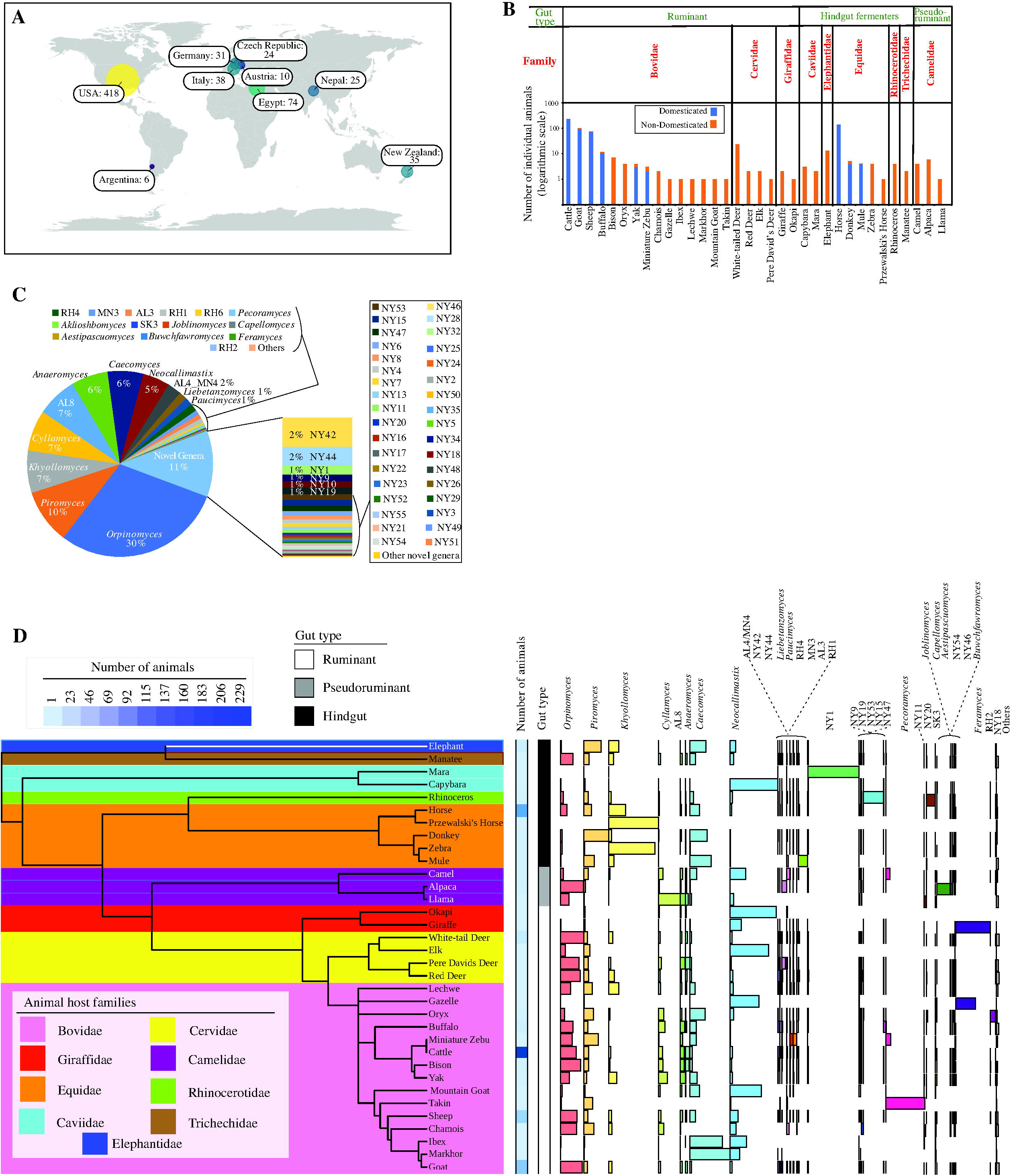
Overview of amplicon datasets analyzed in this study. (A) Map showing the geographical locations and the number of fecal samples analyzed in this study. (B) Stacked bar plot showing the number of samples belonging to each animal species. Animals are ordered by their gut type, then by the animal family. Domestication status is color-coded (domesticated, blue; non-domesticated, orange). (C) Pie chart showing the total percentage abundance of various AGF genera identified in the entire 8.73 million sequence dataset. The relative abundances of the novel genera delineated in this study are shown as a stacked column on the right. Genera whose abundance never exceeded 1% in any of the samples are collectively designated as “others”. (D) AGF community composition by animal species. The phylogenetic tree showing the relationship between animals was downloaded from timetree.org. The tracks to the right of the tree depict the number of individuals belonging to each animal species (shown as a heatmap), and the gut type (color coded as follows: Ruminants, white; Pseudoruminants, grey; Hindgut fermenters, black). AGF community composition for each animal species is shown to the right.

A total of 8.73 million Illumina sequences of the hypervariable region 2 of the large ribosomal subunit (D2 LSU) were obtained. Rarefaction curve (Fig. S1) and coverage estimates (Table S3) demonstrated that the absolute majority of genus-level diversity was captured. The overall composition of the dataset showed a high genus-level phylogenetic diversity, with representatives of 19 out of the 20, and 10 out of the 11, currently described genera, and yet-uncultured candidate genera, respectively, identified (Fig. 1c, d, S2, Table S3). Ubiquity (number of samples in which a taxon is identified) and relative abundance (percentage of sequences belonging to a specific taxon) of different genera were largely correlated (R^2^=0.71, Fig. S3).

To confirm that these patterns were not a function of the primer pair, or sequencing technology (Illumina) employed, we assessed the reproducibility of the observed patterns by conducting a parallel sequencing effort on a subset of 61 samples using a different set of primers targeting the entire D1/D2 LSU region (∼700 bp D1/D2), and a different sequencing technology (SMRT PacBio). A highly similar community composition was observed when comparing datasets generated from the same sample using Illumina versus SMRT technologies, as evident by small Euclidean distances on CCA ordination plot between each pair of Illumina versus PacBio sequenced sample (Fig. S4b-d)), Ordination-based community structure analysis indicated that the sequencing method employed had no significant effect on the AGF community structure (Canonical correspondence analysis ANOVA *p-value*=0.305) (Fig. S4).

### Expanding Neocallimastigomycota diversity

Interestingly, 996,374 sequences (11.4% of the total) were not affiliated with any of the 20 currently recognized AGF genera or 11 candidate genera. Detailed phylogenetic analysis grouped these unaffiliated sequences into 56 novel genera, designated NY1-NY56 (Fig. 2a, Table S3), hence expanding AGF genus-level diversity by a factor of 2.75. In general, relative abundance of sequences affiliated with novel genera was higher in ruminants (Wilcoxon test adjusted p-value <2×10^−16^), as well as in pseudoruminants (Wilcoxon test adjusted p-value =0.02) compared to hindgut fermenters (Fig. 2b-d, Table S4). On the other hand, there was no significant difference in relative abundance of novel genera based on domestication status (Wilcoxon test adjusted p-value =0.69) (Fig. 2e, Table S4).

**Fig. 2.**
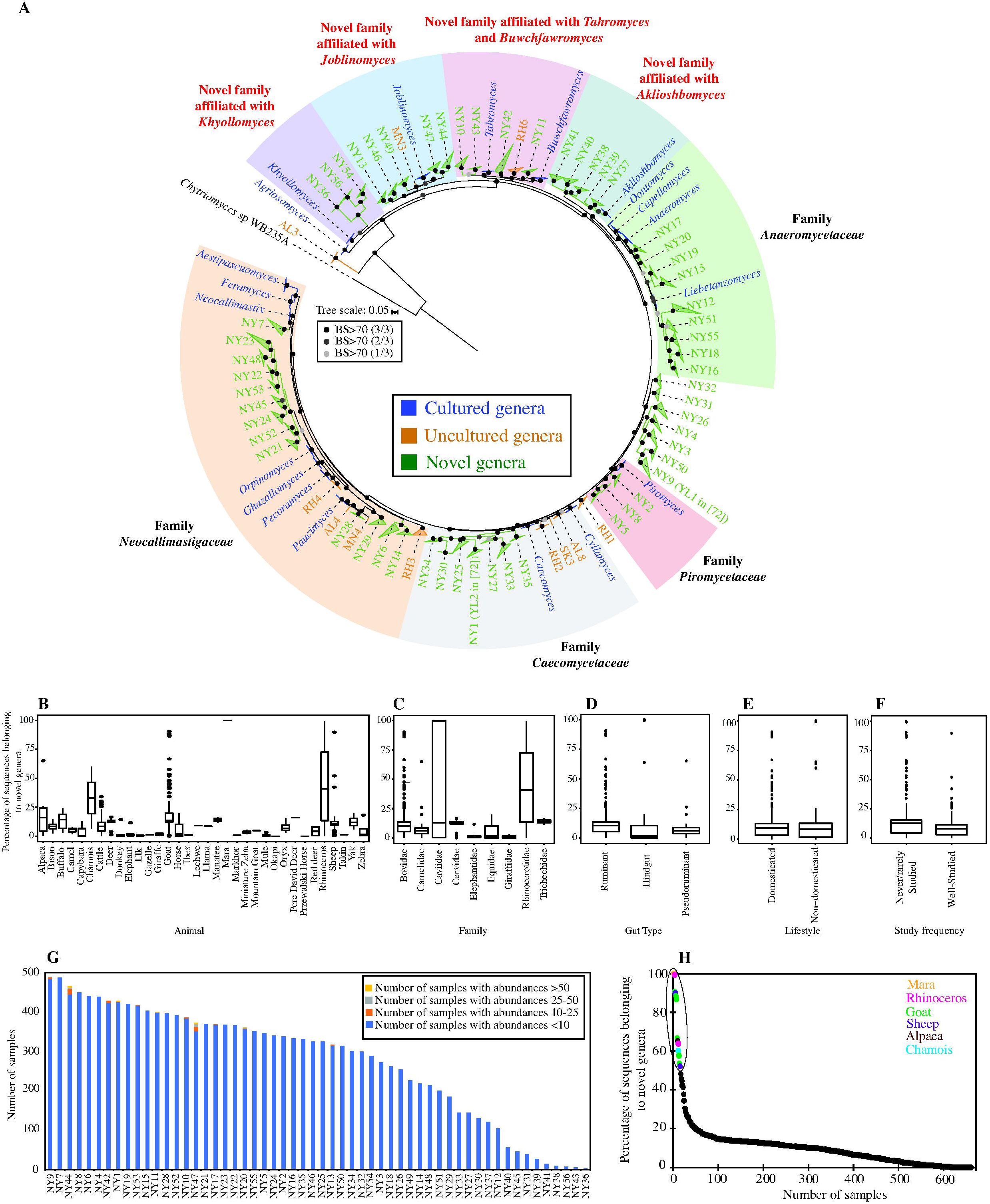
Expanding Neocallimastigomycota diversity. (A) Maximum likelihood phylogenetic tree highlighting the position of novel AGF genera (NY1-NY56, green) identified in this study. The tree includes representatives from all previously reported cultured (blue), and uncultured genera (orange) as references. Two of the 56 novel genera identified here correspond to two novel clades identified in a recent publication: NY1 corresponds to Neocallimastigaceae clade YL2, and NY9 corresponds to Neocallimastigaceae clade YL1 in [72], and both names are acknowledged in the figure. Putative affiliations of novel identified genera with existing AGF families, affiliation with orphan genera, or position as completely novel families are highlighted. The three bootstrap support values (SH-aLRT, aBayes, and UFB) are shown as colored dots as follows: all three support values >70%, black dot; 2/3 support values >70%, dark grey; 1/3 support values >70%, light grey. (B-F) Variation in the proportion of sequences affiliated with novel genera between different animal species (B), animal families (C), animal gut type (D), domestication status (E), and study frequency (F). The distribution of the percentage of novel genera is shown as box and whisker plots, and the results of Wilcoxon test of significance are shown in Table S4. (G-H) Distribution patterns of novel AGF genera identified in this study. (G) Ubiquity of novel genera in analyzed samples. (H) Percentage of sequences belonging to novel genera in each of the 661 samples. The 16 samples that harbored a community with >50% novel sequences are highlighted and color-coded by the animal species as shown in the key.

A closer look at the patterns of distribution of novel genera identified three important trends. First, the proportion of sequences belonging to novel genera in previously well-sampled animals (cattle, sheep, goats, horses, and donkey) was significantly smaller (Wilcoxon test adjusted p-value =2.3×10^−10^) (Fig. 2f, Table S4) than in rarely sampled or previously unsampled hosts (e.g. buffalo, bison, deer, elephant, mara, capybara, manatee, among others), highlighting the importance of sampling hitherto unsampled or rarely sampled animals as a yet-unexplored reservoir for AGF diversity. Second, some novel genera were extremely rare and often identified solely in few sample replicates of a well-sampled animal (e.g. NY42, NY9, NY53, and NY17, in only 5, 2, 1, and 1 cattle samples, respectively), highlighting the importance of replicate sampling for accurate assessment of hosts’ novel pangenomic diversity (Fig. 2g). Finally, 5 of the 56 novel genera were never identified in > 0.1% abundance in any sample, and 16 of the 56 never exceeded 1%, a pattern that highlights the value of deep sequencing to access perpetually rare members of the AGF community (Fig. 2h, Table S5).

Phylogenetically, 32 of the novel lineages identified clustered within the 4 recently proposed families in the Neocallimastigomycota [41], with 13, 7, 9, and 3 genera clustering with the families *Neocallimastigaceae, Caecomycetaceae, Anaeromycetaceae*, and *Piromycetaceae*, respectively). Another 17 novel genera formed additional 4 well-supported family-level clusters with orphan cultured genera (5, 4, 5, and 3 novel genera forming family-level clusters with the orphan genera *Joblinomyces, Buwchfawromyces-Tahromyces, Aklioshbomyces*, and *Khoyollomyces*, respectively). The remaining 7 novel genera were not affiliated with known cultured or uncultured genera and potentially formed novel family-level lineage(s) within the Neocallimastigomycota (Fig. 2a).

Confirmation of the occurrence of such an unexpectedly large number of novel AGF genera and simultaneous recovery of full-length sequence representatives (∼700 bp covering the D1/D2 regions) was achieved by examining the SMRT-PacBio output generated from a subset (61 samples) of the total dataset, as described above. A total of 49 of the 56 novel genera were identified in the PacBio dataset (Table S6). No additional new genera were found using this supplementary sequencing approach. Further, comparing SMRT-versus Illumina-generated tree topologies, revealed nearly identical topologies, phylogenetic distinction, and putative family-level assignments for all novel genera identified (Fig. S5, Table S7).

### Stochastic and deterministic processes play an important role in shaping AGF community

Normalized stochasticity ratios (NST) calculated based on two β-diversity indices (abundance-based Bray-Curtis index, and incidence-based Jaccard index) suggested that both stochastic and deterministic processes contribute to shaping AGF community assembly (Figure 3a-h, Table S8). However, significant differences in the relative importance of these processes were observed across datasets regardless of the β-diversity index used. Specifically, hindgut fermenters and pseudoruminants exhibited significantly lower levels of stochasticity when compared to ruminants (Fig. 3a, e, Wilcoxon adjusted p-value <2×10^−16^). This was also reflected at the animal family level (Fig. 3b, f), as well as at the animal species level (Fig. 3c, g). On the other hand, NST values were highly similar for domesticated versus non-domesticated animals (Fig. 3d, h). To further quantify the contribution of specific deterministic (homogenous and heterogenous selection) and stochastic (homogenizing dispersal, dispersal limitation, and drift) processes in shaping the AGF community assembly, we used a two-step null-model-based quantitative framework that makes use of both taxonomic (RC_Bray_) and phylogenetic (βNRI) β-diversity metrics [33, 34]. Results (Fig. 3i) confirmed a lower overall level of stochasticity in hindgut fermenters, similar to the patterns observed using NST values. More importantly, the results indicate that homogenous selection (i.e., selection of specific taxa based on distinct difference between examined niches) represents the sole (99.8%) deterministic process shaping community assembly across all datasets (Fig. 3i). Within stochastic processes, drift played the most important role in shaping community assembly (83.4% of all stochastic processes), followed by homogenizing dispersal (16.6% of all stochastic processes), with negligible contribution of dispersal limitation. As such, homogenous selection, drift and homogenizing dispersal collectively represented the absolute (>99%) drivers of AGF community assembly, albeit with different relative importance of the three processes in datasets belonging to different animal species, family, gut type, or lifestyle (Fig. 3i).

**Fig. 3.**
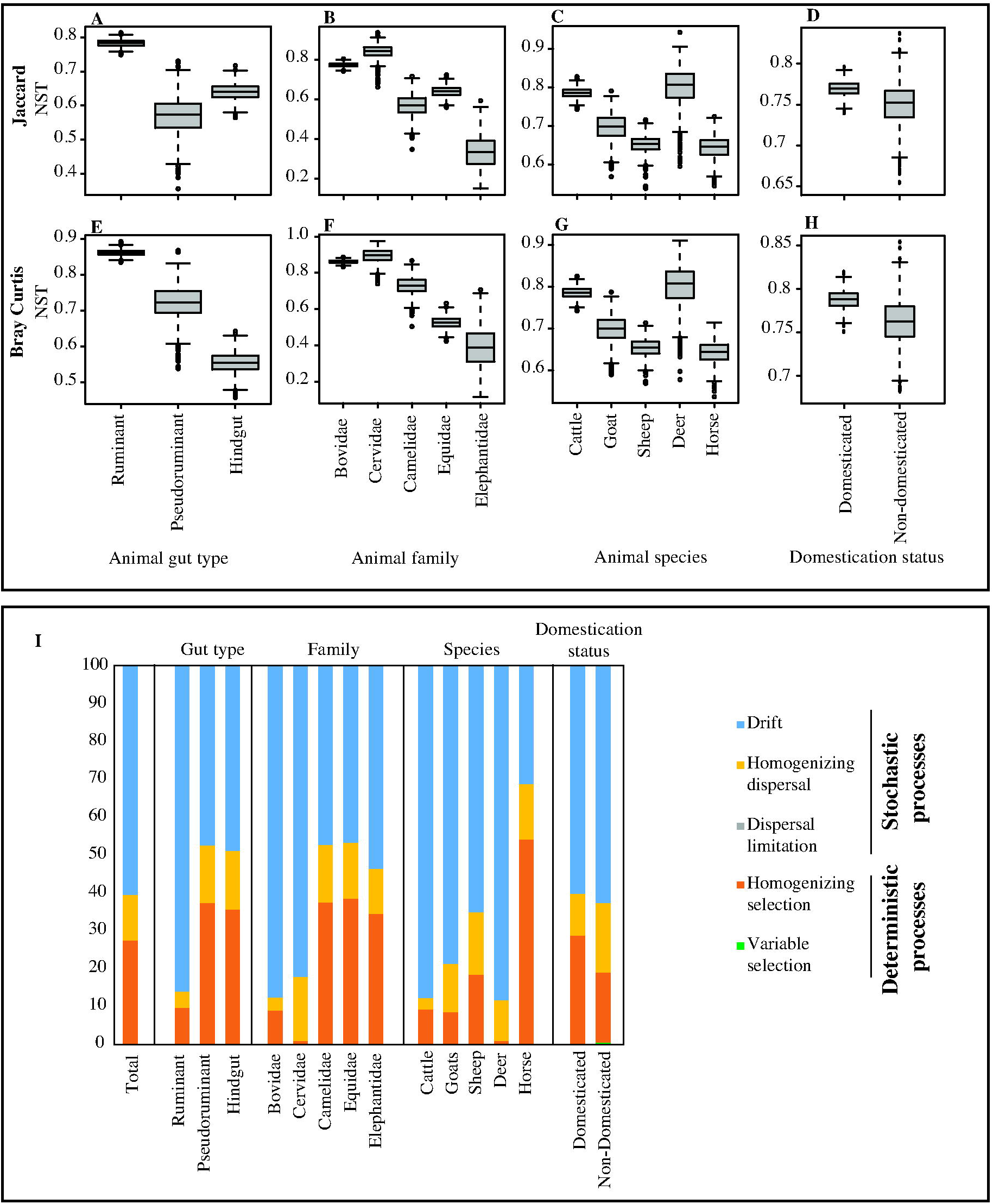
Contribution of stochastic and deterministic processes to AGF community assembly. (A-H) Levels of stochasticity in AGF community assembly were compared between different gut types (A, E), animal families (B, F; for families with more than 10 individuals), animal species (C, G; for animals with more than 20 individuals), and animal domestication status (D, H). Two normalized stochasticity ratio (NST) were calculated; the incidence-based Jaccard index (A-D), and the abundance-based Bray-Curtis index (E-H). The box and whisker plots show the distribution of the bootstrapping results (n=1000). (I) The percentages of the various deterministic and stochastic processes shaping AGF community assembly of the total dataset, and when sub-setting for different animal gut types, animal families, animal species, and animal lifestyles.

### Community structure analysis reveals a strong pattern of fungal-host phylosymbiosis

Assessment of alpha diversity patterns indicated that gut type, animal family, animal species, but not domestication status, significantly affected alpha diversity (Fig S6). Hindgut fermenters harbored a significantly less diverse community compared to ruminants. Within ruminants, no significant differences in alpha diversity levels were observed across various families (Cervidae and Bovidae) or species (deer, goat, cattle, and sheep) (Fig. S6).

Patterns of AGF community structure were assessed using ordination plots (PCoA, NMDS, and RDA) constructed using dissimilarity matrix-based (Bray-Curtis) and phylogenetic similarity-based (unweighted and weighted Unifrac) beta diversity indices (PCoA, and NMDS), or genera abundance data (RDA). The results demonstrated that host-associated factors (gut type, animal family, animal species) play a more important role in shaping the AGF community structure when compared to domestication status, with samples broadly clustering by the animal species (Fig. 4a). PERMANOVA results demonstrated that, regardless of the beta diversity measure, all factors significantly explained diversity (F statistic p-value=0.001), with animal species explaining the most variance (14.7-21 % depending on the index used), followed by animal family (5.4-7.2 %), and animal gut type (4 -5.4 %). Host domestication status only explained 0.4-0.5 % of variance and was not found to be significant with unweighted Unifrac (F statistic p-value =0.143) (Fig. 4b).

**Fig. 4.**
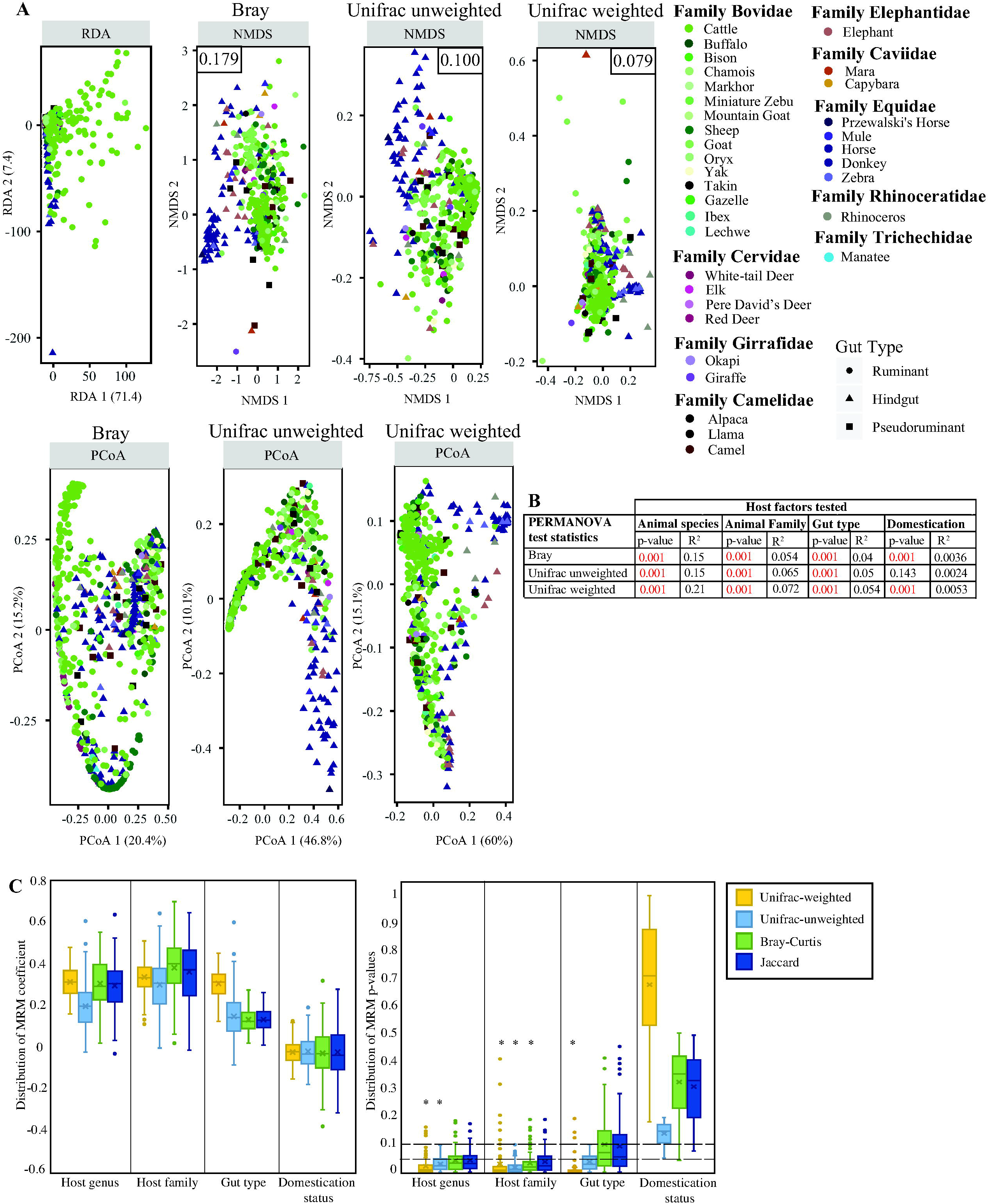
Patterns of AGF beta diversity. (A) Ordination plots based on AGF community structure in the 661 samples studied here. RDA plot was constructed using the genera abundance data. Non-metric dimensional scaling (NMDS) plots were based on both dissimilarity matrix-based (Bray-Curtis), as well as phylogenetic similarity-based (unweighted and weighted Unifrac) indices as shown above each plot. NMDS stress value is shown in the upper corner of each plot. Principal coordinate analysis (PCoA) plots were based on the same three beta diversity measures as shown above each plot, and the % variance explained by the first two axes are displayed on the axes. Samples are color coded by animal species, while the shape depicts the gut type. (B) Results of PERMANOVA test for partitioning the dissimilarity among the sources of variation (including animal species, animal family, animal gut type, and animal lifestyle) for each of the three beta diversity measures used. The F statistic p-value depicts the significance of the host factor in affecting the community structure, while the PERMANOVA statistic R^2^ depicts the fraction of variance explained by each factor. (C) Results of MRM analysis permutation (100 times, where one individual per animal species was randomly selected). Box and whisker plots are shown for the distribution of both the MRM coefficients (left) and the corresponding p-values (right) for the 100 permutations for each of the host factors (animal species, animal family, animal gut type, and animal lifestyle) and dissimilarity indices used (Unifrac weighted, Unifrac unweighted, Bray-Curtis, and Jaccard). If the p-value□was significant (<□0.05) in 75 or more permutations, the host factor was considered significantly affecting community structure (shown as an asterisk above the box and whisker plot). These results were significant for some of the indices used (both Unifrac measures for animal species, both Unifrac measures and Bray-Curtis for animal family, and only weighted Unifrac for animal gut type). Domestication status showed low R^2^ regression coefficients and was found to be not significant using any of the four dissimilarity indices.

Due to the inherent sensitivity of PERMANOVA to heterogeneity of variance among groups [42], we used three additional matrix comparison-based methods: multiple regression of matrices (MRM), Mantel tests for matrices correlations, and Procrustes rotation [43, 44], to confirm the role of host-related factors in shaping AGF community. Results of matrices correlation using each of the three methods, and regardless of the index used, confirmed the importance of animal host species, family, and gut type in explaining the AGF community structure (Fig. S7). Further, we permuted the MRM analysis (100 times), where one individual per animal species was randomly selected for each permutation. Permutation analysis (Fig. 4c) yielded similar results to those obtained from the entire dataset (Fig. S7b), demonstrating that the obtained results are not affected by community composition variation among hosts of the same animal species.

Collectively, our results suggest a pattern of phylosymbiosis, with closely related host species harboring similar AGF communities [45]. To confirm the significant association between the host animal and the AGF community, we employed PACo (Procrustes Application to Cophylogenetic) analysis with subsampling one individual per host species (n=100 subsamples), and compared the distribution of PACo Procrustes residuals of the sum of squared differences within and between animal species (Fig. 5a), animal families (Fig. 5b), and animal gut types (Fig. 5c). Within each animal species, family, or gut type, the variation in PACo residuals were minimal, where 90% of the residuals within animal species ranged from 0.0056 (buffalo) to 0.029 (elephant), within animal family ranged between 0.0048 (Giraffidae) to 0.029 (Elephantidae), and within gut type ranged between 0.007 (foregut) to 0.051 (hindgut). On the other hand, PACo residuals differed significantly between datasets (Wilcoxon two-sided adjusted p-value < 0.01) when animals belonged to different families, or different gut types were examined (Fig. 5a-c, Table S9). These results indicated a strong cophylogenetic signal that was robust to intra-animal species microbiome variation.

**Fig. 5.**
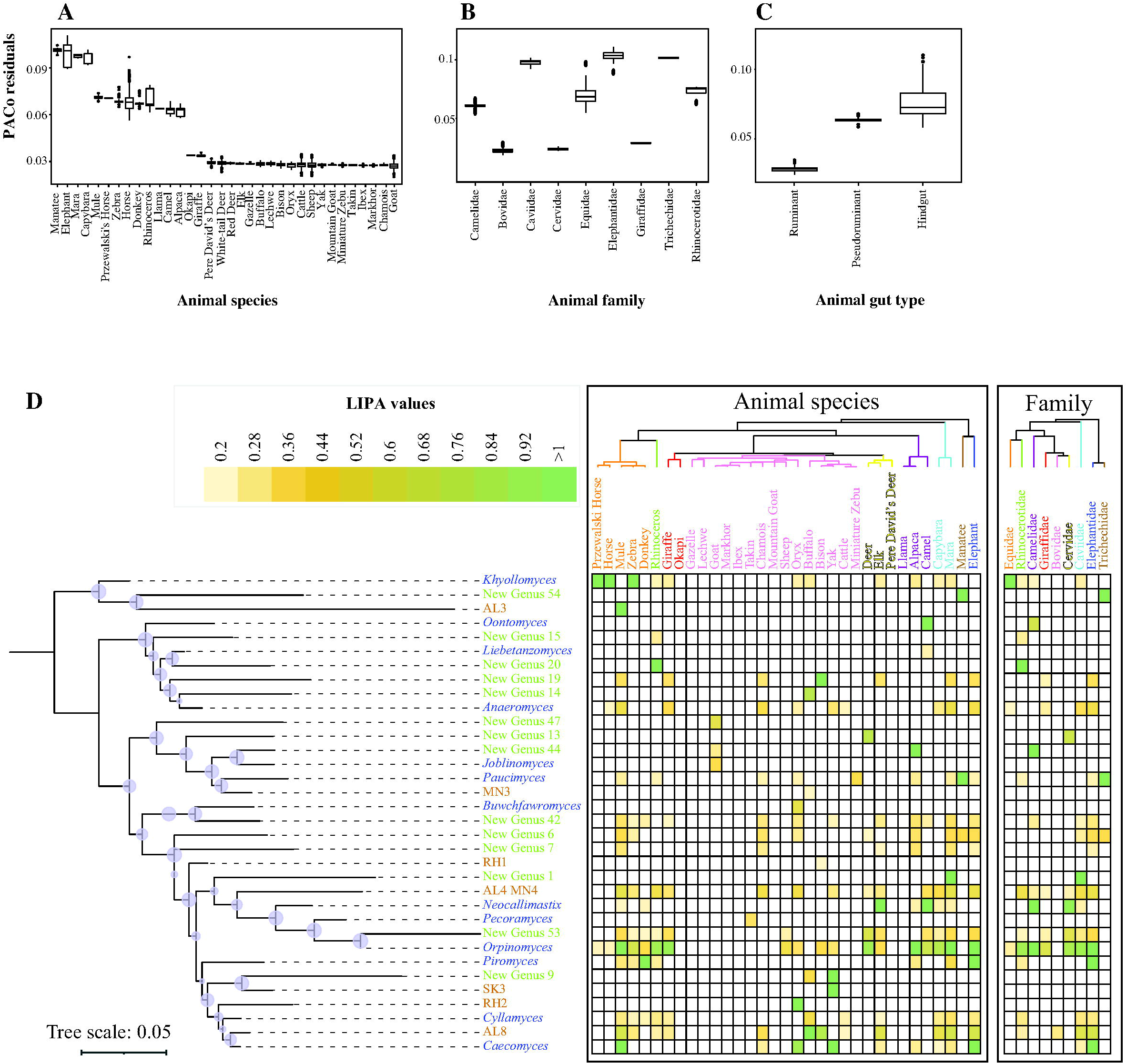
Phylosymbiosis patterns assessed using Procrustes Application to Cophylogenetic (PACo) analysis and Local Indicator of Phylogenetic Association (LIPA). To test the robustness of the phylogenetic signal of association between host phylogeny and the AGF community, PACo analysis was repeated 100 times while subsampling one individual per host genus. The box and whisker plots show the distribution of PACo Procrustes residuals of the sum of squared differences within different animal species (A), animal families (B), and animal gut types (C). Results of two-sided Wilcoxon test for significance of difference between PACo residuals are shown in Table S9. (D) Local indicator of phylogenetic association (LIPA) values for correlations between genera abundances and specific hosts. The AGF tree on the left is a maximum likelihood mid-point rooted tree including only the 34 genera that were found to have significant associations with at least one animal host (LIPA values ≥ 0.2, p-value<0.05). Bootstrap support is shown (as purple circles) for nodes with >70% support. Average LIPA values for specific AGF genus-host genus association (left) and AGF genus-host family association (right) are shown as a heatmap. The host animal tree and host family tree on top were downloaded from timetree.org. Animals are color coded by their respective family and colors follow the same scheme as in Fig. 1d.

### Identifying specific genus-host associations

Global phylogenetic signal statistics (Abouheif’s Cmean, Moran’s I, and Pagel’s Lambda) identified 37 genera with significant correlations to the host phylogenetic tree (p-value < 0.05 with at least one statistic) (Table S10). In addition to global phylogenetic signal statistics, we calculated local indicator of phylogenetic association (LIPA) values for correlations between specific genera abundances and specific hosts. Of the above 37 genera, 34 showed significant associations with at least one animal host (LIPA values ≥ 0.2), with 17 showing strong associations (LIPA values ≥ 1) with specific animal species, and 10 showing strong associations (LIPA values ≥ 1) with certain animal families (Fig. 5d). A distinct pattern of strength of association was observed: All hindgut fermenters exhibited a strong association with a few AGF genera: Horses, Przewalski’s horses, and zebras with the genus *Khoyollomyces*, mules with the uncultured genus AL3, *Orpinomyces*, and *Caecomyces*, donkeys with *Piromyces*, elephants with *Piromyces, Caecomyces*, and *Orpinomyces*, rhinoceros with NY20, manatees with NY54 and *Paucimyces*, and mara with NY1 and *Orpinomyces*. Members of the animal family Equidae mostly showed association with the phylogenetically related genera *Khoyollomyces* and the uncultured genus AL3, suggesting a broader family-level association between both host and fungal families (Fig. 5d, Table S11). On the other hand, a much smaller number of strong host-AGF associations were observed in ruminants (5/22 animal species: NY19 in bison, RH2 in oryx, AL8 in buffalo, NY9, SK3, and *Caecomyces* in yak, and *Neocallimastix* in elk) (Fig. 5d, Table S11). However, this lack of strong LIPA signal was countered by the identification of multiple intermediate and weak cophylogenetic signals (LIPA values 0.2-1, yellow in Fig. 5d) per animal species. It therefore appears that an ensemble of genera, rather than a single genus, is mostly responsible for the phylosymbiosis signal observed in ruminants.

Indeed, DPCoA ordination biplot showed a clear separation of the hindgut families Equidae, and Rhinocerotidae, from the ruminant families Bovidae, Cervidae, and Giraffidae, with the pseudoruminant family Camelidae occupying an intermediate position. This confirmed the patterns suggested by LIPA values, with 14 genera contributing to the foregut community as opposed to only 9 for hindgut fermenters (Fig. S8).

### Phylogenomic and molecular clock analyses correlate fungal-host preferences to co-evolutionary dynamics

The observed patterns of fungal-animal host preferences could reflect co-evolutionary symbiosis (i.e., a deep, intimate co-evolutionary process between animal hosts and AGF taxa). Alternatively, the observed preferences could represent a post-evolutionary environmental filtering process, where prevalent differences in *in-situ* conditions (e.g., pH, retention time, redox potential, feed chemistry) select for adapted taxa from the environment regardless of the partners’ evolutionary history [46]. To address both possibilities, we generated new transcriptomic datasets for 20 AGF strains representing 13 genera, and combined these with 32 previously published AGF transcriptomes [35-40]. We then used the expanded dataset (52 taxa, 14 genera) to resolve the evolutionary history of various AGF genera and estimate their divergence time. In general, most genera with preference to hindgut fermenters occupied an early-diverging position in the Neocallimastigomycota tree, and a broad concordance between their estimated divergence estimate and that of their preferred host family was observed. The genus *Khoyollomyces*, showing preference to horses and zebras (family Equidae), represented the deepest and earliest branching Neocallimastigomycota lineage, with a divergence time estimate of 67-50 Mya (Fig. 6). Such estimate is in agreement with that of the Equidae ∼56 Mya [47, 48]. As well, while the genera AL3 and NY54 are uncultured, and hence not included in the timing analysis, their well-supported association with *Khoyollomyces* in LSU trees (Fig. 2a and S5) strongly suggests a similar early divergent origin. This is in agreement with the early evolution of the families of their hindgut preferred hosts: mules (family Equidae) for AL3, and manatee (family Trichechidae, evolving ∼55 Mya [48]) for NY54). Similarly, the genus *Piromyces*, with a preference to elephants (family Elephantidae) and donkeys (Equidae), also evolved early (55-41 Mya), in accordance with the divergence time estimates for families Equidae and Elephantidae (∼55 Mya) [47, 48]. Finally, the early divergence time estimate of *Paucimyces* (50-38 Mya) is again in agreement with its preference to the hindgut family Trichechidae (Manatee) [48].

**Fig. 6.**
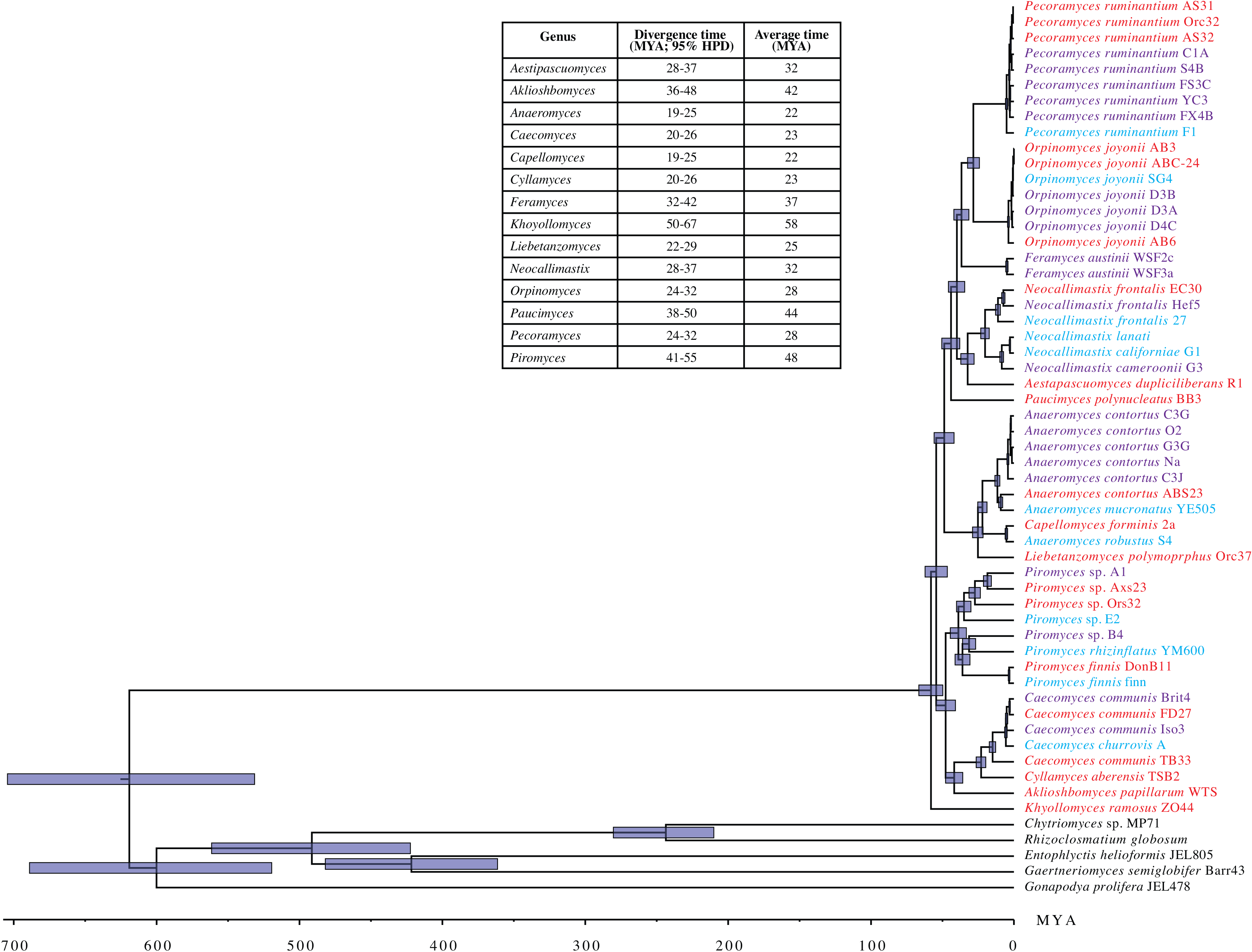
Bayesian phylogenomic maximum clade credibility (MCC) tree of Neocallimastigomycota with estimated divergence time. The isolate names are color coded to show data produced in this study (red), in previous studies by our group (purple) [35, 36] and by other groups (cyan) [37-40, 81, 82]. All clades above the rank of the genus are fully supported by Bayesian posterior probabilities. The 95% highest-probability density (HPD) ranges (blue bars) are denoted on the nodes. For clarity, the average divergence time and 95% HPD ranges of each genus are summarized in a side table.

Contrasting with the basal origins of AGF genera associated with hindgut fermenters, the majority of AGF genera showing strong, intermediate, or weak association with ruminants appear to have more recent evolutionary divergence time estimates. These include many of the currently most abundant and ecologically successful genera, e.g. *Orpinomyces* (24-32 Mya), *Neocallimastix* (28-37 Mya), *Anaeromyces* (19-25 Mya), and *Cyllamyces* (20-26 Mya). Such timing is in agreement with estimates for the rapid diversification and evolution of the foregut fermenting high ruminant families Bovidae, Cervidae, and Giraffidae (18-23 Mya) [30, 49], following the establishment and enlargement of the functional rumen [30].

While these results suggest the central role played by co-evolutionary phylosymbiosis in shaping AGF community, timing estimates for a few genera showed a clear discourse between evolutionary history and current distribution patterns. Such discourse suggests a time-agnostic post-evolutionary environmental filtering process. The late-evolving genera *Orpinomyces* (24-32 Mya) and *Caecomyces* (20-26 Mya) were widely distributed and demonstrated intermediate and strong preferences not only to ruminants, but also to hindgut fermenters (Fig. 5d), suggesting their capacity to colonize hindgut-fermenting hosts, the existence of which has preceded their own evolution. Collectively, these results argue for a major role for co-evolutionary phylosymbiosis and a minor role for post-evolutionary environmental filtering in shaping the AGF community in mammals.

## Discussion

Global amplicon-based, genomic, and metagenomic catalogs have significantly broadened our understanding of microbial diversity on Earth [50-54]. In this study, we generated and analyzed a global (661 samples, 34 animal species, 9 countries, and 6 continents) LSU amplicon dataset, as well as a comprehensive transcriptomic dataset (52 strains from 14 genera) for the Neocallimastigomycota. We focused on using this dataset for documenting the global scope of AGF diversity, as well as deciphering patterns and determinants of the herbivorous mycobiome. However, the size, coverage, and breadth of both datasets render them valuable resources for addressing additional questions and hypotheses by the scientific community.

Our study demonstrates that the scope of AGF diversity is much broader than previously suggested from prior efforts [26, 55, 56]. We identified 56 novel AGF genera, greatly expanding the reported AGF genus-level diversity (Fig. 2a). This broad expansion could be attributed to at least three factors: First, we examined previously unsampled and rarely sampled hosts, including manatee (a herbivorous marine mammal), mara, capybara, chamois, markhor, and takin. Indeed, a greater proportion of sequences belonging to novel genera were found in such samples (Fig. 2), and hence we posit that examining the yet-unsampled herbivorous mammals should be prioritized for novel AGF discovery. Second, we examined a large number of replicates per animal species, and found that some novel genera were detected in some but not all samples from the same animal. Given the immense number of herbivores roaming the Earth, it is rational to anticipate that additional AGF diversity surveys of even well sampled hosts could continue to yield additional novel lineages. Third, we accessed rare members of the AGF community through deep sequencing, and found that 5 of the 56 novel genera were never identified in > 0.1% abundance in any sample, and 16 of the 56 never exceeded 1% (Table S5). The rationale for the existence, maintenance, and putative ecological role of rare members within a specific ecosystem has been highly debated [57]. We put forth two distinct, but not mutually exclusive, explanations for the maintenance of rare AGF taxa. First, rare taxa could persist in nature by coupling slow growth rates to superior survival (e.g., high oxygen tolerance, formation of resistant structures outside the herbivorous gut), dispersal, and transmission capacities when compared to more abundant taxa. Second, rare taxa could provide valuable ecological services under specific conditions not adequately captured by the current sampling scheme, e.g. specialization in attacking specific minor components in the animal’s diet, superior growth in specific cases of gut dysbiosis, or during early stages of their hosts’s life. In newborn animals, the undeveloped nature of the alimentary tract [58], the liquid food intake, and distinct behavior, e.g. coprophagy in foals, may select for a distinct microbiome, and rare AGF members of the community could hence represent remnants of the community developing during the early days of the host life. Detailed analysis of the effect of dysbiosis on AGF communities, as well as the temporal development patterns from birth to maturity is needed to experimentally assess the plausibility of both scenarios.

Our results highlight the importance of the hitherto unrecognized role of stochastic processes (drift and homogenizing dispersal) in shaping AGF community in herbivores. The contribution of these processes was on par with (in the hindgut fermenting and pseudoruminant families), or exceeding (in the ruminant families Bovidae and Cervidae) that of deterministic niche-based processes (Fig. 3i). We attribute the high contribution of drift to the restricted habitat and small population sizes of AGF in the herbivorous gut, conditions known to elicit high levels of drift [34]. As well, the highly defined functional role for AGF in the herbivorous gut (initial attack and colonization of plant biomass), high levels of similarity in metabolic abilities, substrates preferences, and physiological optima across genera argue for a null-model scenario, where phylogenetically distinct taxa roles are ecologically interchangeable. The importance of homogenizing dispersal (Fig. 3i) suggests a high and efficient dispersal rate leading to community homogenization. While the strict anaerobic nature of AGF could argue that dispersal limitation, rather than homogenizing dispersal, should be more important in shaping AGF community. However, such a perceived transmission barrier is readily surmounted via direct vertical mother-to-offspring transfer by post-birth grooming, as well as direct horizontal transmission between animals, or through feed-fecal cross contamination in close quarters [59].

A greater level of stochasticity was observed in ruminants compared to hindgut fermenters. This could be due to the proximity of the prominent AGF-harboring chamber (rumen) to the site of entry (mouth) in ruminants, compared to the distant location of the reciprocal chamber (caecum) in hindgut fermenters. This results in a greater rate of secondary airborne transmission in foregut fermenters, as well as a greater level of selection for AGF inoculum in hindgut fermenters during their passage through the alimentary tracts (with various lengths and resident times). As well, the observed pattern could be due to the high-density rearing conditions and higher level of inter-species cohabitation between many ruminants (e.g. cattle, sheep, goats), as opposed to the relatively lower density and cross-species cohabitation for hindgut fermenters (e.g. horses, elephants, manatees).

While stochastic processes play a role in AGF community assembly, the role of deterministic processes remains substantial (Fig. 3). Host-associated factors are logical factor to examine as key drivers of AGF community structure. Differences in overall architecture, size, and residence time in alimentary tracts of different hosts could result in niche-driven selection of distinct AGF communities. In addition, variation in bacterial and archaeal community structures between hosts could also elicit various levels of synergistic, antagonistic, or mutualistic relationships that impact AGF community. However, domestication status could counter, modulate, or override host identity. Domesticated animals are fed regularly and frequently a monotonous diet, compared to the more sporadic feeding frequency and more diverse feed types experienced by non-domesticated animals. Such differences could select for AGF strains suited for each lifestyle. Furthermore, the close physical proximity and high density of animals in domesticated settings are conducive to secondary airborne transmission, while the more dispersed lifestyle of wild herbivores could elicit a more stable community within a single animal species.

Ordination clustering patterns and PERMANOVA analysis demonstrated that host-associated factors explained a much higher proportion of the observed variance, when compared to host’s domestication status (Fig. 4b). Such relative importance of host-associated factors was further confirmed by multivariate regression approaches (multiple regression of matrices, Mantel tests for matrices correlations, and Procrustes rotation Table S9, Fig. 4C). Global phylogenetic signal statistics (Table S10), LIPA (Fig. 5d, Table S11), and PACo analysis (Fig. 5, S7) confirmed this observed cophylogenetic pattern and identified and quantified the strength of AGF-host associations. All hindgut fermenters exhibited strong associations with a few AGF genera, while multiple intermediate cophylogenetic signals were identified for foregut fermenters. This suggests that enrichments of an ensemble of multiple genera, rather than a single genus, is mostly responsible for the distinct community structure observed in foregut fermenters. These patterns of strong animal-host correlation are in agreement with the patterns of lower stochasticity (Fig. 3), and lower alpha diversity (Fig. S6) observed in hindgut fermenters.

As described above, the predicted role of phylosymbiosis in shaping AGF community structure in extant animal hosts could reflect two distinct, but not mutually exclusive, mechanisms; co-evolutionary phylosymbiosis, and post-evolutionary host filtering. Documenting such a relationship between hosts and their microbiomes requires a strong backbone of evolutionary trees where phylogenies are accurately resolved, and evolutionary timing is well described. While this has been achieved for mammalian hosts [60], the phylogenetic and evolutionary relationships between various genera within the Neocallimastigomycota are less certain, with topologies recovered from single locus phylogenetic analyses often dependent on the locus examined, region amplified, taxa included in the analysis, and tree-building algorithm employed [26, 27, 61]. Phylogenomic approaches using whole genomic and/or transcriptomic datasets are a promising tool for resolving such relationships [62-66]. Our results from transcriptomics-enabled phylogenomic and molecular clock analysis strongly suggest a more prevalent role for co-evolutionary phylosymbiosis in shaping the observed pattern of AGF diversity. Specifically, it appears that the evolution of various herbivorous mammalian families, genera, and species following the K-Pg extinction event and continuing through the early Miocene, and the associated evolutionary innovations in alimentary tract architecture (e.g. evolution of the three-chambered forestomach of pseudoruminants, and the four-chambered stomach of ruminants), drove a parallel evolutionary diversification process within the Neocallimastigomycota. This is supported by the preference of earliest divergent AGF genera to hindgut fermenting hosts, e.g. *Khoyollomyces* and associated genera (AL3 and NY54) to members of the Equidae [6, 67, 68], as well as the general basal position of additional hindgut-preferring genera, e.g. *Piromyces* (41-55 Mya) and *Paucimyces* (38-50 Mya). This is in agreement with the fact that early mammals roaming the Earth past the K-Pg boundary (∼65.5 Mya) were hindgut fermenters. On the other hand, the recent origin for the foregut-preferring genera *Orpinomyces, Neocallimastix*, and *Anaeromyces* (22-32 Mya) suggests this followed the earlier evolution (∼ 40 Mya) of a functional and enlarged rumen [30], and the subsequent rapid diversification and evolution of multiple families in the high ruminants (Suborder Ruminantia, Infraorder Pecora), e.g. Bovidae, Cervidae, Giraffidae (18-23 Mya) [30, 49]. As such, organismal and gut evolution appear to have provided novel niches that triggered rapid AGF genus-level diversification in the early Miocene. However, in addition to phylosymbiosis, post-evolutionary host filtering also appears to play a role in shaping the AGF community. For example, members of the genus *Orpinomyces* showed a strong association to a wide range of animal families and gut types (Fig. 6). The reason for the ecological success of *Orpinomyces* in multiple hosts is currently uncertain, but members of this genus exhibit robust polycentric growth pattern, enabling fast vegetative production via hyphal growth and fragmentation.

In addition to host phylogeny, and domestication status, additional factors could impact AGF community structure. These factors include biogeography, animal age, animal sex, as well as diet. However, the effect of these non-host-related factors on community structure could potentially be conflated when examined across different hosts. One way to avoid such conflation is to limit the analysis to the same animal species (e.g. examining the effect of biogeography on the AGF community structure using cattle samples only). Our analyses suggest a possible role for such factors in shaping AGF community structure across a single species (Fig. S9). The country of origin significantly explaining 3.9% of variance in cattle (F-test p-value=0.002), 5.6% of variances in horses (F-test p-value=0.012), 23.6% of variances in goats (F-test p-value=0.001), and 30.8% of variances in sheep (F-test p-value=0.001). Similarly, animal age significantly explained 3.5-32.7% (depending on the animal species), and animal sex significantly explained 2.1-15.0% (depending on the animal species) of variances in AGF community structure. As such, while these results suggest a putative role for such factors in shaping AGF diversity, future controlled studies are needed to examine each issue while normalizing others e.g. sampling cattle of the same breed, at the same age, and feeding regime but housed in different geographic locations to examine biogeographic patterns).

In summary, our results demonstrate that the scope of fungal diversity in the herbivorous gut is much broader than previously implied from prior culture-dependent, culture-independent, and –omics surveys [26, 38, 69-71], quantify the relative contribution of various ecological factors in shaping AGF community assembly across various hosts, and demonstrate that host-specific evolutionary processes (e.g. evolution of host families, genera, and gut architecture) played a key role in driving a parallel process of AGF evolution and diversification.

## Materials and Methods

### Sampling and DNA extraction

A total of 661 fecal samples belonging to 34 different mammalian animal species and 9 families of ruminant, pseudoruminant, and hindgut fermenters were included in the final analysis (Fig. 1a-b, Table S2). Samples were obtained from 15 different research groups using a single standardized procedure (Supp. Methods). DNA extractions were conducted in eight laboratories using DNeasy Plant Pro Kit (Qiagen®, Germantown, Maryland) according to manufacturer’s instructions.

### Illumina sequencing

All PCR amplification reactions, amplicon clean-up, quantification, index and adaptor ligation, and pooling were conducted in a single laboratory (Oklahoma State University, Stillwater, OK, USA) to eliminate inter-laboratory variability. All reactions utilized the DreamTaq Green PCR Master Mix (ThermoFisher, Waltham, Massachusetts), and AGF-LSU-EnvS primer pair (AGF-LSU-EnvS For: 5’-GCGTTTRRCACCASTGTTGTT-3’, AGF-LSU-EnvS Rev: 5’-GTCAACATCCTAAGYGTAGGTA-3’) [72] targeting a ∼370 bp region of the LSU rRNA gene (corresponding to the D2 domain), an amplicon size enabling high throughput sequencing using the Illumina MiSeq platform. Pooled libraries (300-350 samples) were sequenced at the University of Oklahoma Clinical Genomics Facility (Oklahoma City, Oklahoma) using the Illumina MiSeq platform (Supp. methods).

### Complementary PacBio sequencing

As a complimentary approach to Illumina sequencing, we conducted PacBio sequencing on a subset (n=61) of the Illumina-sequenced samples to amplify the D1/D2 LSU region (∼700 bp). Primers utilized were the fungal forward primer (NL1: 5’-GCATATCAATAAGCGGAGGAAAAG-3’), and the AGF-specific reverse primer (GG-NL4: 5’-TCAACATCCTAAGCGTAGGTA-3’) [26, 73]. Details on the rationale for PacBio sequencing, as well as PCR amplification, amplicon clean-up, quantification, index and adaptor ligation, and pooling are in the Supp. methods.

### Sequence processing, and taxonomic and phylogenetic assignments

Protocols for read assembly, and sequence quality trimming, as well as procedures for calculating thresholds for species and genus delineation and genus-level assignments are provided in Supp. methods. Briefly, pairwise sequence divergence estimates comparison between SMRT and Illumina amplicons showed very high correlation (R^2^= 0.885, Fig. S10), and indicated that the 2% sequence divergence cutoff previously proposed as the threshold for delineating AGF species using the D1/D2 region (based on comparisons of validly described species) [28] is equivalent to 3.5% using the D2 region only, and the 3% sequence divergence cutoff previously proposed as the threshold for delineating AGF genera using the D1/D2 region [28] is equivalent to 5.1% using the D2 region only (Fig. S10). Assignment of sequences to AGF genera was conducted using a two-tier approach for genus-level phylogenetic placement as described previously [26, 28] and as detailed in the Supp. Methods.

### Role of stochastic versus deterministic processes in shaping AGF community assembly

We assessed the contribution of various deterministic and stochastic processes to the AGF community assembly using both normalized stochasticity ratio (NST) [32], and the null-model-based quantitative framework implemented by [33, 34]. The NST ratio infers ecological stochasticity, however, values do not pinpoint the sources of selection (determinism) or stochasticity. Also, NST values are calculated solely based on taxonomic diversity indices with no consideration to the phylogenetic turnover in the community. To quantify the contribution of various deterministic (homogenous and heterogenous selection) and stochastic (dispersal preference, limitation, drift) processes in shaping the AGF community assembly, we used a two-step null-model-based quantitative framework that makes use of both taxonomic (RC_Bray_) and phylogenetic (βNRI) β-diversity metrics [33, 34] (Supp. methods).

### Factors impacting AGF diversity and community structure

We considered two types of factors that could potentially impact AGF diversity and community structure: host-associated factors, and non-host-associated factors. For host-associated factors, we considered animal species, animal family, and animal gut type, while for non-host-associated factors we considered domestication status, biogeography (country of origin), age, and sex. For testing the effect of biogeography, age, and sex, we carried out comparisons only on samples belonging to the same animal species in an attempt to control for other host-associated factors. For these comparisons, only the four mostly sampled animal species (cattle, goats, sheep, and horses) were considered.

Alpha diversity estimates were calculated as described in the supplementary document. All beta diversity indices (both dissimilarity matrix-based e.g., Bray-Curtis, as well as phylogenetic similarity-based e.g., unweighted and weighted Unifrac were calculated using the ordinate command in the Phyloseq R package. The pairwise values were used to construct ordination plots (both PCoA and NMDS) using the function plot_ordination in the Phyloseq R package. RDA plots were also constructed using the genera abundance data. To partition the dissimilarity among the sources of variation (including animal host species, animal host family, animal gut type, and animal lifestyle), PERMANOVA tests were run for each of the above beta diversity measures using the vegan command Adonis, and the F-statistics p-values were compared to identify the host factors that significantly affect the AGF community structure. The percentage variance explained by each factor was calculated as the percentage of the sum of squares of each factor to the total sum of squares.

To further quantitatively assess factors that explain AGF diversity, we used three additional multivariate regression approaches based on matrices comparison: multiple regression of matrices (MRM), Mantel tests for matrices correlations, and Procrustes rotation. Bray-Curtis, Jaccard dissimilarity, Unifrac weighted, and Unifrac unweighted dissimilarity matrices were compared to a matrix of each of the host factors tested (animal host species, animal host family, animal gut type, and animal lifestyle) using the commands MRM, and mantel in the ecodist R package, for running multiple regression on matrices, and Mantel tests. The Procrustes rotation was calculated using the protest command in the vegan R package. The significance, and importance of the host factor in explaining the AGF community structure were assessed by comparing the p-values, and coefficients (R^2^ regression coefficients of the MRM analysis, Spearman correlation coefficients of the Mantel test, and symmetric orthogonal Procrustes statistic of the Procrustes analysis), respectively. Finally, to assess the sensitivity of multivariate regression methods to community composition variation among hosts of the same species, we permuted the MRM analysis 100 times, where one individual per animal species was randomly selected. For each of these permutations, and for each dissimilarity matrix-host factor comparison, a p-value and an R^2^ regression coefficient was obtained. We considered a host factor significant in explaining AGF community structure, if in the permutation analysis the p-value□ obtained was significant (p <□0.05) in at least 75 permutations (Supp. methods).

### Assessing phylosymbiosis patterns

To test for patterns of phylosymbiosis, and the presence of a cophylogenetic signal between the animal host and the AGF genera constituting the gut community, we used Procrustes Application to Cophylogenetic Analysis (PACo) through the paco R package (Supp. methods). For pinpointing specific animal host-fungal associations, we employed two approaches. We first used the phyloSignal command in the phylosignal R package to calculate three global phylogenetic signal statistics, Abouheif’s Cmean, Moran’s I, and Pagel’s Lambda. The values of these statistics plus the associated p-values were employed to identify the AGF genera that have a significant association with an animal host. We then used the lipaMoran command in the phylosignal R package to calculate LIPA (Local Indicator of Phylogenetic Association) values for each sample-AGF genus pair, along with the associated p-values of association significance. For AGF genera showing significant associations (LIPA p-values < 0.05), we calculated average LIPA values for each animal host species, and animal family. We considered average LIPA values in the range of 0.2-0.4 to represent weak associations, in the range 0.4-1 to represent moderate associations, and above 1 to represent strong associations. To further explore the notion that enrichments of an ensemble of multiple genera, rather than a single genus, is responsible for the distinct community structure observed in ruminants and pseudoruminants, we used the ordinate command in Phyloseq followed by plot_ordination to construct a double principal coordinate analysis (DPCoA) plot.

### Transcriptomic analysis

Prior studies by our research group have generated 21 transcriptomes from 7 genera [35, 36]. Here, we added 20 transcriptomes from 7 additional genera, isolated during a long term multi-year isolation effort in the authors laboratory [26, 28], and included an extra 11 publicly available transcriptomic datasets [37-40]. The dataset of 52 transcriptomes was used for phylogenomic analysis as described in [41]. For RNA extraction, cultures grown in rumen fluid-cellobiose medium [74] were vacuum filtered then grounded with a pestle under liquid nitrogen. Total RNA was extracted using Epicentre MasterPure yeast RNA purification kit (Epicentre, Madison, WI) according to manufacturer’s instructions. Transcriptomic sequencing using Illumina HiSeq2500 platform and 2□×□150 bp paired-end library was conducted using the services of a commercial provider (Novogene Corporation, Beijing, China), or at the Oklahoma State University Genomics and Proteomics center. The RNA-seq data were qualitytrimmed and *de novo* assembled with Trinity (v2.6.6) using default parameters. Redundant transcripts were clustered using CD-HIT [75] with identity parameter of 95% (–c 0.95), and subsequently used for peptide and coding sequence prediction using the TransDecoder (v5.0.2) (https://github.com/TransDecoder/TransDecoder) with a minimum peptide length of 100 amino acids. BUSCO [76] was used to assess transcriptome completeness using the fungi_odb10 dataset modified to remove 155 mitochondrial protein families as previously suggested [37]. In addition, five Chytridiomycota Genomes (*Chytriomyces* sp. strain MP 71, *Entophlyctis helioformis* JEL805, *Gaertneriomyces semiglobifer* Barr 43, *Gonapodya prolifera* JEL478, and *Rhizoclosmatium globosum* JEL800) were included to provide calibration points. The same phylogenomic dataset (670 protein-coding genes) produced for [41] was used as the original input. Gap regions were removed using trimAl v1.4 [77]. Alignment files that contained no missing taxa and were longer than 150 nucleotide sites were selected for subsequent analyses. By employing a greedy search in PartitionFinder v2.1.1 [78], the 88 selected alignments were grouped into 15 partitions with independent substitution models. All partition files and respective models were loaded in BEAUti v1.10.4 [79] with calibration priors specified as previously described [36] ((i) a direct fossil record of Chytridiomycota from the Rhynie Chert (407 Mya) & (ii) the emergence time of Chytridiomycota (573 to 770 Mya as 95% HPD)) for Bayesian inference and divergence time estimation implemented in BEAST v1.10.4. The Birth-Death incomplete sampling tree model was employed for interspecies relationship analyses. Unlinked strict clock models were used for each partition independently. Three independent runs were performed for 50 million generations and Tracer v1.7.1 [80] was used to confirm that sufficient effective sample size (ESS>200) was reached after the default burn-in (10%). The maximum clade credibility (MCC) tree was compiled using TreeAnnotator v1.10.4 [79].

### Sequence and data deposition

Illumina reads were deposited in GenBank under BioProject accession number PRJNA887424, BioSample accession numbers SAMN31166910-SAMN31167478, and SRA accessions SRR21816543-SRR21817111. PacBio sequences were deposited in GenBank as a Targeted Locus Study project under the accession KFWW00000000. The version described in this paper is KFWW01000000. PacBio sequence representatives of the 49 novel AGF groups were deposited in GenBank under accession numbers OP253711-OP253963 (Table S6). Raw Illumina RNA-seq read sequences are deposited in GenBank under the BioProject accession number PRJNA847081, BioSample accession numbers SAMN28920465-SAMN28920484, and individual SRA accessions SRR19612694-SRR19612713.

## Code availability

Code for phylogenomic analysis (Fig. 6) is available at https://github.com/stajichlab/PHYling_unified. All Code used to create all other figures is available at https://github.com/nohayoussef/AGF_Mammalian_Herbivores.

## Contributions

Conceptualization and experimental design by NHY and MSE, Sample collection, archiving, and laboratory experimentation by HM, ALJ, AXA, JH, CJP, RAH, ASY, MAT, CDM, PHJ, MS, PR, MN, JMV, SMP, ALG, JH, DY, KF, DJG, RV, GM, SM, MTK, YIN, SSD, APF. Data analysis by CHM, YW, JES, JAE, NHY, MSE. Manuscript writing by NHY and MSE. Funding acquisition by CDM, PHJ, GM, MN, JMV, SMP, DY, NHY, and MSE.

## Acknowledgments

We thank Kelle Sundstrom, Emily Looper and Ruth Scimeca (Oklahoma State University College of Veterinary Medicine), Jennifer D’Agostino, and Rebecca Snyder (Oklahoma City Zoo), as well as multiple ranchers and farmers throughout the USA for colleting and providing samples. We also thank David Hume, Katherine Lowe, Richard Muirhead, Christian Sauermann, Shengjing Shi and Gosia Zobel who contributed the range of NZ samples. We also thank ZOO Garden Ostrava, Farm Tehov, and Farm Duběnka Ostrava (Czech Republic) for help with sample collections. We also thank Katelyn Pixley, Ryzzah Trinidad, and Russell Croy for technical Assistance,.

