## Supplementary document for "Patterns and determinants of the global herbivorous mycobiome"

^12^Prague Zoo, Prague, Czechia.

^13^Department of Veterinary Medicine, University of Sassari, Sardinia, Italy.

^14^University of Milan, Dept. of Agricultural and Environmental Sciences, Milan, Italy.

^15^ Anaerobic Fungi Network, Kerkdriel, Netherlands.

^16^Bioenergy Group, Agharkar Research Institute, Pune, India.

^17^ Oklahoma State University, Department of Animal and Food Sciences, Stillwater, Oklahoma, USA.

**I. Supplementary materials and methods**

**Sampling.** A total of 661 samples belonging to thirty-three different mammalian animal species and nine families of foregut ruminant, foregut pseudoruminant, and hindgut fermenters were included in the final analysis (Figures 1a-b, Table S2). Samples were obtained from 15 different laboratories using a standardized procedure. Fresh, moist fecal samples originating from a single animal were scooped immediately post defecation into sterile 50-ml plastic tubes and rapidly sealed. Samples were transferred to the laboratory on ice, usually within one hour, and stored at -20ºC prior to DNA extraction. All individuals collecting samples from domesticated animals were owners or caretakers and were authorized and trained in animal keeping and husbandry. For domesticated animals reared in a research institution, all IRB protocols for animal rearing were observed. Sampling wild herbivores was conducted through a partnership with hunting communities, alleviating the need for obtaining hunting permits. All hunters had the appropriate licenses, and the animals were shot on public land during the hunting season.

**DNA extraction.** DNA extractions were conducted in eight laboratories (Oklahoma, USA: 418 samples, Egypt: 74 samples, Nepal: 25 samples, Austria: 10 samples, Italy:38 Samples, Czechia: 24 samples, Germany: 31 samples, and New Zealand: 35 samples) and followed the manufacturer’s instructions for DNeasy Plant Pro Kit (Qiagen®, Germantown, Maryland). Few samples were from a prior study [1]. The kit was shipped from USA to labs in Nepal and Egypt, and was purchased independently elsewhere.

**PCR amplification, and Illumina sequencing.** The 28S rRNA was used as the phylomarker for this diversity survey since as previously suggested for AGF genus-level delineation [2, 3]. The LSU rRNA has been regarded as a more reliable marker for AGF diversity characterization (compared to the general fungal phylomarker ITS1 region) by the scientific community [4], and has been previously employed in AGF culture-independent diversity surveys [5]. By comparison, the ITS1 region suffers from high within-strain sequence divergence, and length variability [5, 6]. All PCR amplifications were conducted in a single laboratory to eliminate inter-laboratory variability. All reactions utilized the DreamTaq Green PCR Master Mix (ThermoFisher, Waltham, Massachusetts), and primers targeting the D2 region of the LSU rRNA (AGF-LSU-EnvS For: 5’-GCGTTTRRCACCASTGTTGTT-3’, AGF-LSU-EnvS Rev: 5’-GTCAACATCCTAAGYGTAGGTA-3’) for amplification. The primers were recently developed and extensively evaluated for sensitivity, specificity, Neocallimastigomycota coverage, and ability to differentiate between all known AGF genera and candidate genera [7]. The primers target a ~370 bp region of the LSU rRNA gene (corresponding to the D2 domain), hence allowing for high throughput sequencing using the Illumina MiSeq platform. Primers were modified to include the Illumina overhang adaptors. PCR reactions contained 2 µl of DNA, 25 µl of the DreamTaq 2X master mix (Life Technologies, Carlsbad, California), 2 µl of each primer (10 µM) in a 50 µl reaction mix. The PCR protocol consisted of an initial denaturation for 5 min at 95 °C followed by 40 cycles of denaturation at 95 °C for 1 min, annealing at 55 °C for 1 min and elongation at 72 °C for 1 min, and a final extension of 72 °C for 10 min. PCR products were individually cleaned to remove unannealed primers using PureLink® gel extraction kit (Life Technologies, Carlsbad, California), and the clean product was used in a second PCR reaction to attach the dual indices and Illumina sequencing adapters using Nexterra XT index kit v2 (Illumina Inc., San Diego, California). These second PCR products were then cleaned using PureLink® gel extraction kit (Life Technologies, Carlsbad, California), individually quantified using Qubit® (Life Technologies, Carlsbad, California), and pooled using the Illumina library pooling calculator (https://support.illumina.com/help/pooling-calculator/pooling-calculator.htm) to prepare 4-5 nM libraries. Pooled libraries (300-350 samples) were sequenced at the University of Oklahoma Clinical Genomics Facility using the MiSeq platform.

**Complementary PacBio sequencing.** The 700-750 bp D1/D2 LSU fragment has recently been adopted [4] as the gold standard for AGF genus- and species-level assignments, as well as for circumscribing boundaries between various AGF clades. However, the length of the region precludes the use of Illumina technology for high throughput sequencing and necessitates the use of the technically more cumbersome and expensive Sanger or PacBio sequencing platforms. Therefore, as a complimentary approach to Illumina sequencing, we conducted PacBio sequencing on a subset of the Illumina-sequenced samples (n=61) with the following goals in mind: (1) to ensure that community membership, structure, and diversity estimates obtained with the shorter fragment (D2 region only) are comparable to those obtained with the previously utilized D1/D2 region, (2) to ensure the feasibility and resolution power of the D2 region in identifying and differentiating various fungal lineages, and (3) to confirm the presence of the unexpectedly high number of putative novel genera, provide full-length representative sequences [4], and amend the curated D1/D2 LSU rRNA database currently handled by the authors as part of the broader AGF community of researchers ([www.anaerobicfungi.org)](http://www.anaerobicfungi.org)) . For amplification of the D1/D2 LSU region, we paired a universal fungal forward primer (NL1: 5’- GCATATCAATAAGCGGAGGAAAAG-3’) with an AGF-specific reverse primer (GG-NL4: 5’-TCAACATCCTAAGCGTAGGTA-3’) [5, 8]. Primers were barcoded to allow multiplexing and PacBio sequencing. The PCR protocol consisted of an initial denaturation for 5 min at 95 °C followed by 40 cycles of denaturation at 95 °C for 1 min, annealing at 55 °C for 1 min and elongation at 72 °C for 1 min, and a final extension of 72 °C for 10 min. Amplicons were purified using PureLink® gel extraction kit (Life Technologies), quantified using Qubit® (Life Technologies), pooled, and sequenced at Washington State University core facility using one cell of the SMRT Pacific Biosciences (PacBio) Sequel II system.

**Illumina and PacBio sequence processing:** Forward and reverse Illumina reads were assembled using make.contigs command in mothur [9], followed by screening to remove sequences with ambiguous bases, sequences with homopolymer stretches longer than 8 bases, and sequences that were shorter than 200 or longer than 380 bp. For PacBio sequences, raw reads were processed using the official PacBio pipeline (RS_Subreads.1) (<http://files.pacb.com/software/smrtanalysis/2.2.0/doc/smrtportal/help/!SSL!/Webhelp/CS_Prot_RS_Subreads.htm>), and filtered using default settings of the minimum read length, and minimum read quality. Remaining reads were then processed with the PacBio RS_ReadsOfInsert protocol (<http://files.pacb.com/software/smrtanalysis/2.2.0/doc/smrtportal/help/!SSL!/Webhelp/CS_Prot_RS_ReadsOfInsert.htm>) for generating single-molecule consensus reads from the insert template. Circular consensus sequences were further processed in mothur [9] to remove any sequence with average quality score < 25, sequences with ambiguous bases, sequences not containing the correct barcode, sequences with more than 2 bp difference in the primer sequence, and/or sequences with homopolymer stretches longer than 8 bp. To identify any CCS with the primer sequence in the middle, we performed a standalone Blastn-short using the primer sequence as the query, and removed the identified sequences using the remove.seqs command in mothur.

**Taxonomic and phylogenetic assignments.** To examine how taxa delineation cutoffs previously proposed based on the D1/D2 region [4, 5] correlate to those of the shorter Illumina-generated D2 LSU fragments obtained in this study, we conducted preliminary comparison of all possible pairwise sequence divergence values from the alignment of the whole D1/D2 region of 206 reference sequences (available at [www.anaerobicfungi.org](http://www.anaerobicfungi.org)), to those from the truncated alignment covering the D2 region only (corresponding to the region that would be amplified using the AGF-LSU-EnvS primer pair above). Sequence divergence estimates from the two sets of alignments were very well correlated (R^2^= 0.885, Figure S9). However, comparison of pairwise sequence divergence using the whole D1/D2 region versus the D2 region suggests that the 2% sequence divergence cutoff previously proposed as the threshold for delineating AGF species using the D1/D2 region (based on comparisons of validly described species) [4] is equivalent to 3.5% using the D2 region only, and the 3% sequence divergence cutoff previously proposed as the threshold for delineating AGF genera using the D1/D2 region [4] is equivalent to 5.1% using the D2 region only (Figure S9).

Therefore, pairwise distances were used to cluster the sequences into species level OTUs using the proposed sequence threshold of 3.5%. On the other hand, prior research has shown that using specific thresholds for genus-level delineation in AGF is problematic. For example, some genera were found to harbor higher intra-genus D1/D2-LSU region sequence divergence values (e.g. the genus *Piromyces* intra-genus D1/D2-LSU region sequence divergence ranges between 0% and 5.7%), while others diverge by <2% from neighboring genera (e.g. the *Anaeromyces-Liebetanzomyces-Capellomyces-Oontomyces* clade harbors inter-genus D1/D2-LSU region sequence divergence values ranging between 1.8% and 2.5%) [4]. Therefore, we refrained from using a predetermined threshold to assign sequences to AGF genera, and instead used a two-tier approach for genus-level phylogenetic placement. First, sequences were compared by Blastn to the curated D1/D2 LSU rRNA AGF database ([www.anaerobicfungi.org](http://www.anaerobicfungi.org)), and sequences were classified as their first hit taxonomy if the percentage similarity to the first hit was > 96% and the two sequences were aligned over >70% of the query sequence length. For all sequences that could not be confidently assigned to an AGF genus by Blastn, insertion into a reference LSU tree (with representatives from all cultured and uncultured AGF genera and candidate genera) was used to assess novelty. Briefly, unaffiliated sequences (100-200 at a time) were aligned to the reference database using align.seqs in mothur, and the alignment was used to construct maximum likelihood phylogenetic trees in FastTree [10] using the GTR model. Sequences were assigned to a novel genus when they cluster as an independent genus-level clade with high (>70%) bootstrap support in the ML tree. Representatives of novel genera were sequentially added to the reference LSU tree, before processing the next batch of unaffiliated sequences. Intra-genus D2 region sequence divergence for sequences assigned to any novel genus never exceeded 5% (equivalent to 3% D1/D2 sequence divergence). Following the assignment of all sequences to either an existing or a novel genus, final trees with 5-10 representatives of each genus were generated in IQtree [11] using the alignment of the D2 region. ModelFinder [12] through IQtree [11] was used to select the best substitution model (based on the lowest BIC criteria). Maximum likelihood trees were constructed under the predicted best model, with the -alrt 1000, the -bb 1000, and the --abayes options added to the commandline for performing the Shimodaira–Hasegawa approximate likelihood ratio test (SH-aLRT), the ultrafast bootstrap (UFB) [13], and the approximate Bayes test. This resulted in the generation of phylogenetic trees with three support values (SH-aLRT, aBayes, and UFB) on each branch. Phylogenetic analysis as described above resulted in the confident assignment of every single sequence to either an existing cultured, or uncultured genus, or to a novel genus. These genus-level assignments were then used to build a taxonomy file in mothur, which was subsequently used to build a shared file using the mothur commands phylotype and make.shared.

Amplicon sequence variants (ASVs) have recently been gaining popularity and momentum in describing diversity in bacterial [14, 15], archaeal [16], and fungal [17] surveys, a proposition augmented by improved sequence quality and stringent quality control procedures on all sequencing platforms [18-21], we, however, refrained from using ASVs in this study, due to the fact that a significant level of within-strain divergence (ranging between 0.1%-1.9% [5]) is observed in the multiple LSU rRNA gene copies (estimated around 170 per genome). As such, use of ASVs, with its emphasis on exact sequencing identity [18-20] would immensely overestimate AGF diversity and bias community structure estimates.

**Role of stochastic versus deterministic processes in shaping AGF community assembly.** We assessed the contribution of various deterministic and stochastic processes to the AGF community assembly using both normalized stochasticity ratio (NST) [22], and the null-model-based quantitative framework implemented by [23, 24]. The NST index infers ecological stochasticity, however, values do not pinpoint the sources of selection (determinism) or stochasticity. Also, NST values are calculated solely based on taxonomic diversity indices with no consideration to the phylogenetic turnover in the community. To quantify the contribution of various deterministic (homogenous and heterogenous selection) and stochastic (dispersal preference, limitation, drift) processes in shaping the AGF community assembly, we used a two-step null-model-based quantitative framework that makes use of both taxonomic (RC_Bray_) and phylogenetic (βNRI) β-diversity metrics [23, 24]. The NST package in R was used to calculate the normalized stochasticity ratio (NST) based on two taxonomic β-diversity dissimilarity metrics; the incidence-based Jaccard index, and the abundance-based Bray-Curtis index, where an NST value of > 50% indicates a more stochastic assembly, while values <50% indicate a more deterministic assembly. To test the significance of difference between pairs of animal species (for animals with more than 20 individuals; cows, goats, sheep, deer, and horses), animal families (for families with more than 10 individuals; Bovidae, Cervidae, Camelidae, Equidae, and Elephantidae), and animal gut types (foregut, pseudoruminant, and hindgut), we used the function nst.boot in the NST package in R to randomly draw samples within each group followed by bootstrapping of NST values. Obtained values were then compared using Wilcoxon test with Benjamini-Hochberg adjustment. The iCAMP R package was used to calculate values of beta net relatedness index (βNRI), and modified Raup-Crick metric based on Bray Curtis metric (RC_Bray_) using the function bNRIn.p to evaluate the turnover for both phylogenetic, and taxonomic diversity. Values of βNRI were used first to partition selective processes into homogenous (number of pairwise comparisons with βNRI values < -2), and heterogenous selection (number of pairwise comparisons with βNRI values > 2). All other pairwise comparisons (with absolute βNRI values < 2) are considered contributing to stochastic processes (not assigned to selection), and can be further broken down into dispersal and drift based on the taxonomic diversity (values of RC_Bray_). Specifically for these, the number of pairwise comparisons with absolute values of RC_Bray_ < 0.95 are considered contributing to drift, while the number of pairwise comparisons with absolute values of RC_Bray_ > 0.95 are considered contributing to dispersal. This last fraction can be further broken down into homogenizing dispersal (RC_Bray_ values <-0.95), and dispersal limitation (RC_Bray_ values >0.95). The contribution of each of these processes (homogenous selection, heterogenous selection, homogenizing dispersal, dispersal limitation, and drift) to the total AGF community assembly was calculated from the corresponding number of pairwise comparisons falling into each category as a percentage of all pairwise comparisons.

**Factors impacting AGF diversity and community structure.** We considered two types of factors that could potentially impact AGF diversity and community structure: host-associated factors, and non-host-associated factors. For host-associated factors, we considered animal species, animal family, and animal gut type, while for non-host-associated factors, we considered animal domestication status, biogeography (country of origin), animal age, and animal sex. For testing the effect of biogeography, age, and sex on alpha diversity measures and community structure, we opted to carry out comparisons only on samples belonging to the same animal species in an attempt to control for other host-associated factors that might conflate the results. For these comparisons, only the four most-sampled animal species (cattle, goats, sheep, and horses) were considered.

**Alpha diversity measures.**

Alpha diversity estimates (observed number of genera, Shannon, Simpson, and Inverse Simpson diversity indices) were calculated using the command estimate_richness in the Phyloseq R package. For comparison of alpha diversity between samples, patterns were assessed in samples with at least 1000 sequences (n=421 samples) using the four indices, and two sampling strategies (with and without random subsampling of 1000 sequences) for eight total comparisons. The importance of various factors (host-associated factors, e.g. gut type, animal family, or animal species; domestication status, and biogeography) in shaping the observed patterns of alpha diversity was examined using ANOVA (calculated using the aov command in R). Only samples that have at least 10 replicates (at any of these host factor levels) were included in the analysis. These included foregut and hindgut (for the gut type factor comparison), families Bovidae, Cervidae, and Equidae (for the animal family comparison), cows, goats, sheep, deer, and horses (for the animal genus comparison), and domesticated and non-domesticated (for domestication status comparison). Additionally, post hoc Tukey HSD tests for multiple comparisons of means were run on the results of ANOVA (using TukeyHSD command in R) for all possible pairwise comparisons to identify the pairs of groups that are significantly different for each host factor.

As mentioned above, we opted to carry out comparisons of the effect of biogeography, age, and sex only on samples belonging to the same animal species (only the four most-sampled animals were included) in an attempt to control for other host-associated factors that might conflate the results. Biogeography comparisons were conducted on cattle, goats, sheep, and horses datasets originating from USA, Egypt, Germany, Italy, Austria, Czech Republic, New Zealand, and Argentina. ANOVA (calculated using the aov command in R) was used to identify the animal species whose AGF alpha diversity significantly differed between countries. For these animal datasets, post hoc Tukey HSD tests for multiple comparisons of means were run on the results of ANOVA (using TukeyHSD command in R) for all possible pairwise country comparisons to identify the pairs that are significantly different for each animal genus. Additionally, the effect of the US state of origin on AGF alpha diversity in cattle and horses was also tested using ANOVA followed by post hoc Tukey HSD tests for multiple comparisons of means for all possible pairwise state comparisons to identify the pairs that are significantly different for each animal species.

Age (young, < 1 year; adult, >1 year), and sex (male versus female) comparisons were also considered only for the four most-sampled animals (cattle, goats, sheep, and horses). ANOVA followed by post hoc Tukey HSD tests were used.

**AGF community structure.** The genus-level shared file was used to calculate several beta diversity indices (including both dissimilarity matrix-based (e.g. Bray-Curtis), as well as phylogenetic similarity-based (e.g. unweighted and weighted Unifrac) using the ordinate command in the Phyloseq R package. The pairwise values were used to construct ordination plots (both PCoA and NMDS) using the function plot_ordination in the Phyloseq R package. RDA plots were also constructed using the genera abundance data. To assess the variability in community structure between samples belonging to each animal host species (only for animals with 4 or more individuals), animal host family, animal gut type, and animal domestication status, we first calculated group centroids for each of these groups using the vegan command betadisper. Following, the ordination distance of each sample to its group centroid was calculated (as the Euclidean distance between two points), and distances from group centroids were plotted in a box and whisker plot (using the command boxplot in R). To partition the dissimilarity among the sources of variation (including animal host species, animal host family, animal gut type, and domestication stauts), PERMANOVA tests were run for each of the above beta diversity measures using the vegan command Adonis, and the F-statistics p-values were compared to identify the host factors that significantly affect the AGF community structure. The percentage variance explained by each factor was calculated as the percentage of the sum of squares of each factor to the total sum of squares.

Due to the inherent sensitivity of PERMANOVA to the heterogeneity of variance among groups [25], and to further quantitatively assess factors that explain AGF diversity, we used three multivariate regression approaches based on matrices comparison: multiple regression of matrices (MRM), Mantel tests for matrices correlations, and Procrustes rotation. Bray-Curtis, and Jaccard dissimilarity matrices were first calculated from the genus shared file using vegdist command in Vegan. Similarly, Unifrac weighted, and Unifrac unweighted dissimilarity matrices were calculated using the distance command in the Phyloseq package. Each of these four AGF dissimilarity matrices were compared to a matrix of each of the host factors tested (animal host species, animal host family, animal gut type, and domestication status). For the animal host genus, a cophenetic matrix was calculated (using the command cophenetic in the ape R package) based on the newick tree downloaded from timetree.org and modified to include all the samples studied here with very short branch length between samples from the same animal species. For the animal host family, animal gut type, and domestication status, since these were nominal values, matrices were constructed by Gower transformation [26]. Each of the AGF community dissimilarity matrices (n=4) was then correlated to each of the host factor matrices (n=4) using the commands MRM, and mantel in the ecodist R package, for running multiple regression on matrices, and Mantel tests, respectively. The Procrustes rotation was calculated using the protest command in the vegan R package. For each of the host factors tested, 12 total correlations (3 methods x 4 dissimilarity indices) were compared to evaluate the importance of the host factor in explaining the AGF community structure. This was achieved by comparing the p-values for significance of correlation, and coefficients (R^2^ regression coefficients of the MRM analysis, Spearman correlation coefficients of the Mantel test, and symmetric orthogonal Procrustes statistic of the Procrustes analysis) for the importance of the factor in explaining community structure. Finally, to assess the sensitivity of multivariate regression methods to community composition variation among hosts of the same species, we permuted the MRM analysis 100 times, where one individual per animal species was randomly selected. For each of these permutations, and for each dissimilarity matrix-host factor comparison, a p-value and an R^2^ regression coefficient is obtained. We considered a host factor significant in explaining AGF community structure, if in the permutation analysis the p-value obtained was significant (p < 0.05) in at least 75 permutations.

To test for the effect of biogeography, sex, and age on the AGF community structure, and to overcome compounded effects from other host factors, we selected a subset of the whole dataset to include only samples from the four most-sampled animals (namely, cattle, goats, sheep, and horses). For each of these animal species, we calculated Bray-Curtis dissimilarity indices using the ordinate command in the Phyloseq R package. The pairwise values were used to construct PCoA ordination plots using the function plot_ordination in the Phyloseq R package. The samples were color coded by country, sex, or age. To test for the significance of the above three factors in describing AGF community structure in each animal genus, PERMANOVA tests were run using the vegan command Adonis. The F-statistics p-value was used to assess the significance of AGF community difference between countries, young versus adult animals, and males versus females, and the sum of squares was used to assess the percentage variance explained by the country of origin for each of the four animal species.

**Assessing phylosymbiosis patterns.** To test for patterns of phylosymbiosis, and the presence of a cophylogenetic signal between the animal host and the AGF genera constituting the gut community, we used Procrustes Application to Cophylogenetic Analysis (PACo) through the paco R package. Briefly, the analysis involves the host cophenetic distance matrix (reflecting the phylogenetic relationships between hosts), the AGF cophenetic distance matrix based on the phylogenetic relationship of the different AGF genera to each other, and the AGF genera abundance in the samples. With these three inputs, the analysis then translates the distance matrices of the animal host and the AGF phylogenies into principal coordinates, followed by rotating one set of the coordinates to maximize superimposition on the other. The sum of squared residuals of this superimposition is calculated and is used as an indication of congruency between the two sets, with smaller sum of squared residuals indicating better congruency. A bias-correction step is also added. The analysis produces, besides the bias-corrected sum of squared residuals, a p-value for the goodness of fit between the two phylogenies. Additionally, to assess the sensitivity of PACo analysis to community composition variation among hosts of the same species, we repeated the analysis while subsampling one individual per host genus (n=100 subsamples), and compared the distribution of PACo Procrustes residuals of the sum of squared differences between different animal species, different animal families, and different gut types. To test for the significance of the difference between residuals, we used Wilcoxon test with Benjamini-Hochberg adjustment.

For pinpointing specific animal host-fungal associations, we employed two approaches. We first used the phyloSignal command in the phylosignal R package to calculate three global phylogenetic signal statistics, Abouheif’s Cmean, Moran’s I, and Pagel’s Lambda. The values of these statistics plus the associated p-values identify the AGF genera that have a significant association with an animal host. We considered any genus with p-value < 0.05 with at least one statistic to be significantly correlated to the host phylogenetic tree. We then used the lipaMoran command in the phylosignal R package to calculate LIPA (Local Indicator of Phylogenetic Association) values for each sample-AGF genus pair, along with the associated p-values of association significance. For AGF genera showing significant associations (LIPA p-values < 0.05), we calculated average LIPA values for each animal host species, and animal family. We considered average LIPA values in the range of 0.2-0.4 to represent weak associations, in the range 0.4-1 to represent moderate associations, and above 1 to represent strong associations.

To further explore the notion that enrichments of an ensemble of multiple genera, rather than a single genus, is responsible for the distinct community structure observed in foregut fermenters, we constructed a double principal coordinate analysis (DPCoA) ordination using the genera with abundance in the top 25%. The genera (n=28) were first selected using the filterfun_sample(topp(0.25)) command in Phyloseq. The DPCoA was constructed using the ordinate command in Phyloseq followed by plot_ordination. DPCoA uses both abundance and phylogenetic information about the samples, allowing both the samples and the taxa to be plotted on the same coordinate space, and thus the Euclidean distance between samples or their group centroids and AGF genera could be compared. Thus, AGF genera with Euclidean distances close to group centroids are considered to contribute more to the community structure of the group. We used betadisper in the R package Vegan to calculate centroids for the three different gut types, the nine different animal families, and the animal genera with at least 4 individuals (n=15), and ggplot2 to draw 95% confidence level ellipses for the three gut types.

**Transcriptomic analysis.** Prior studies by our research group have generated 21 transcriptomes from 7 genera [27, 28]. Here, we added 20 transcriptomes from 7 additional genera, isolated during a long term multi-year isolation effort in the authors laboratory [4, 29] and included an extra 11 publicly available transcriptomic datasets [30-33]. Cultures grown in rumen fluid-cellobiose medium [34] were vacuum filtered then grounded with a pestle under liquid nitrogen. Total RNA was extracted using Epicentre MasterPure yeast RNA purification kit (Epicentre, Madison, WI) according to manufacturer’s instructions. Transcriptomic sequencing using Illumina HiSeq2500 platform and 2 × 150 bp paired-end library was conducted using the services of a commercial provider (Novogene Corporation, Beijing, China), or at the Oklahoma State University Genomics and Proteomics center. The RNA-seq data were quality trimmed and *de novo* assembled with Trinity (v2.6.6) using default parameters. Redundant transcripts were clustered using CD-HIT [35] with identity parameter of 95% (–c 0.95), and subsequently used for peptide and coding sequence prediction using the TransDecoder (v5.0.2) (<https://github.com/TransDecoder/TransDecoder>) with a minimum peptide length of 100 amino acids. BUSCO [36] was used to assess transcriptome completeness using the fungi_odb10 dataset modified to remove 155 mitochondrial protein families as previously suggested [30]. The dataset of 52 transcriptomes was used for phylogenomic analysis as described in [37]. In addition, five Chytridiomycota Genomes (*Chytriomyces* sp. strain MP 71, *Entophlyctis helioformis* JEL805, *Gaertneriomyces semiglobifer* Barr 43, *Gonapodya prolifera* JEL478, and *Rhizoclosmatium globosum* JEL800) were included to provide calibration points. The same phylogenomic dataset (670 protein-coding genes) produced for [37] was used as the original input. Gap regions were removed using trimAl v1.4 [38]. Alignment files that contained no missing taxa and were longer than 150 nucleotide sites were selected for subsequent analyses. By employing a greedy search in PartitionFinder v2.1.1 [39], the 88 selected alignments were grouped into 15 partitions with independent substitution models. All partition files and respective models were loaded in BEAUti v1.10.4 [40] with calibration priors specified as previously described [28] ((i) a direct fossil record of Chytridiomycota from the Rhynie Chert (407 Mya) & (ii) the emergence time of Chytridiomycota (573 to 770 Mya as 95% HPD)) for Bayesian inference and divergence time estimation implemented in BEAST v1.10.4. The Birth-Death incomplete sampling tree model was employed for interspecies relationship analyses. Unlinked strict clock models were used for each partition independently. Three independent runs were performed for 50 million generations and Tracer v1.7.1 [41] was used to confirm that sufficient effective sample size (ESS>200) was reached after the default burn-in (10%). The maximum clade credibility (MCC) tree was compiled using TreeAnnotator v1.10.4 [40].

**II. Supplementary figures.**

**Figure S1: Rarefaction curve** Rarefaction curves showing the increase in the number of genera observed as the number of sequences increase per sample. Rarefaction data were calculated and plotted in R using rarecurve in Vegan.

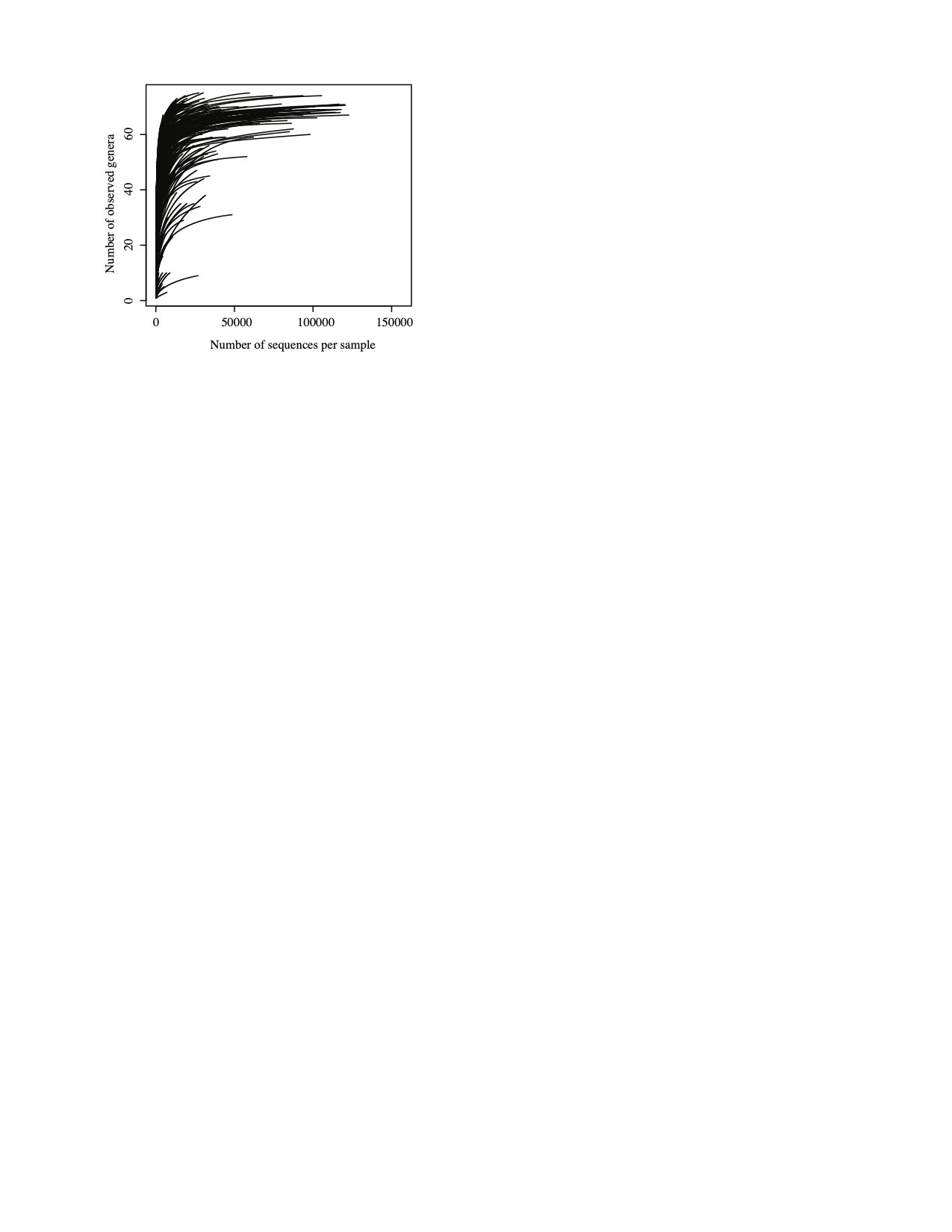

**Figure S2:** **Occurrence and relative abundance distribution of all AGF genera encountered in this study.** (A) Number of samples (out of 661) in which each AGF genus was identified. Genera are shown in descending order of their average percentage abundance across samples. (B) Box and whisker plot showing the distribution of the percentage abundance of the 84 genera. Genera are shown in the same order as in (A).

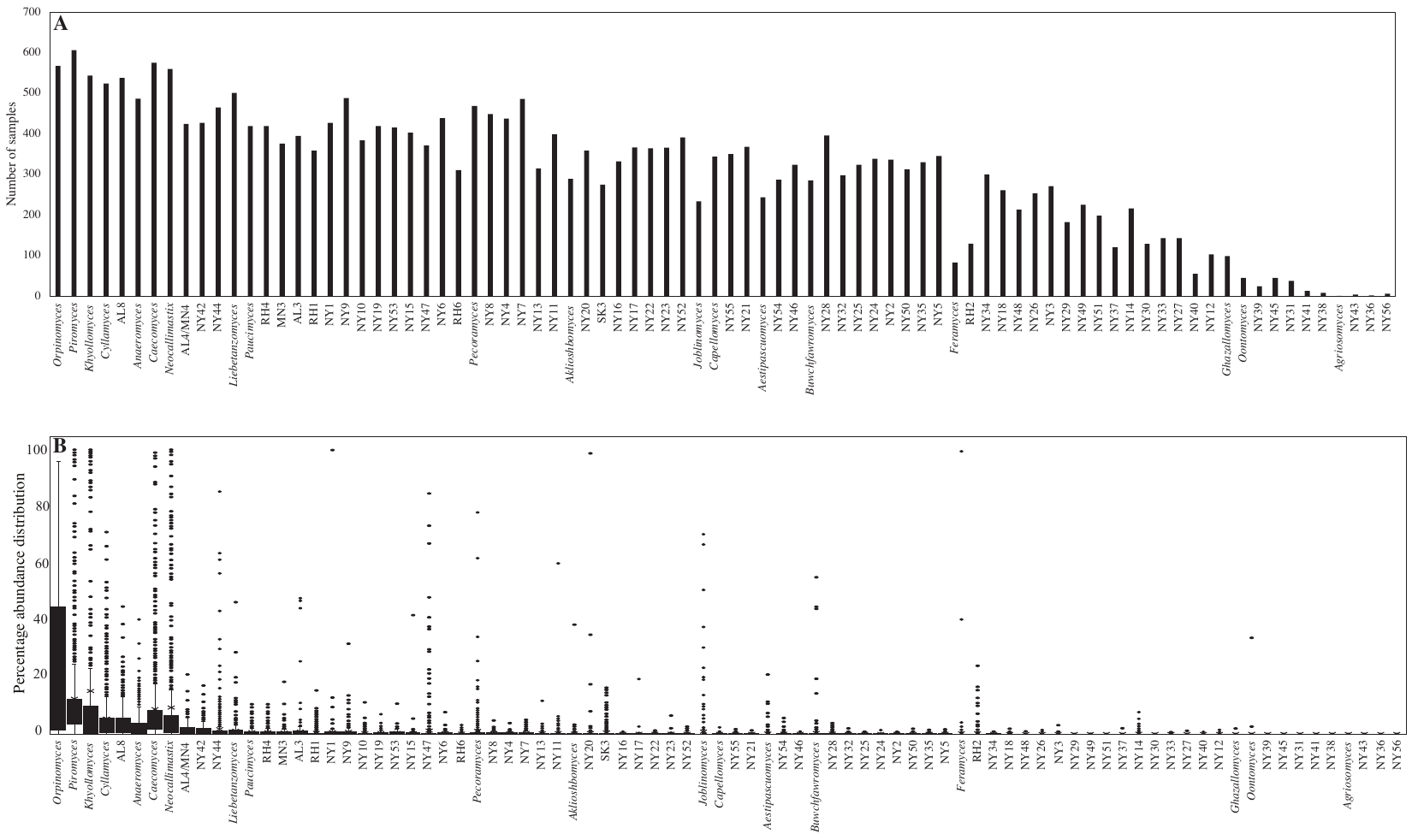

**Figure S3: Abundance-occurrence plots**. Relationship between occurrence (number of samples) and average relative abundance of each of the 84 genera encountered in this study. The number of samples in which the genera were identified is shown on the X-axis. Average percentage abundance across samples is plotted on the Y axis in a logarithmic scale to show genera present below 1% abundance. Occurrence and relative abundance of different genera were largely correlated (R^2^=0.71), with the few highlighted exceptions (*Joblinomyces*, *Feramyces*, *Onotomyces*, and RH2).

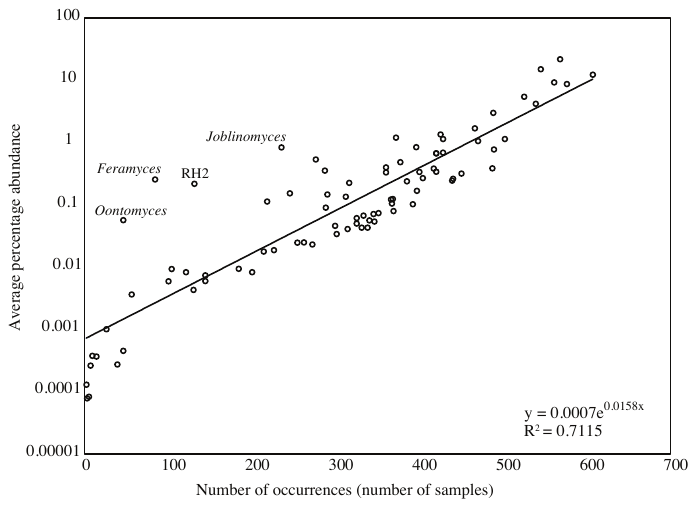

**Figure S4: Comparison of community structure patterns between Illumina- and SMRT PacBio-generated datasets.** Comparative analysis of the community structure and composition in 61 samples (60 cows and 1 bison) that were amplified using the D2-targeting primers and sequenced with Illumina sequencing technology, as well as using a different set of primers targeting the whole D1/D2 region and sequenced using the PacBio sequencing technology.

(A) AGF community composition in the 61 samples sequenced using Illumina (left stacked bar) and SMRT (right stacked bar) sequencing technologies showing the overall similarity in community composition. (B-D) Community structure in the 61 samples sequenced with the two sequencing technologies. (B) Canonical correspondence analysis (CCA) of the AGF community in the 122 samples (61 samples x 2 sequencing technologies) showing the similarity in community between Illumina-sequenced (red) and SMRT-sequenced (blue) samples. The community structures of the same sample sequenced with the two sequencing technologies were similar (clustered close to each other on the CCA plot). This is evident from the comparisons of the distribution of Euclidean distances on the CCA plot between all possible pairs of Illumina-sequenced samples (red), all possible pairs of SMRT-sequenced samples (blue), and the 61 pairs of Illumina versus PacBio sequenced samples (grey) (C), where the Euclidean distances between the Illumina-SMRT pairs are smaller. Data for the grey box and whisker plot in (C) (the 61 Illumina-SMRT pairs) is shown in detail in (D). Fifty-two pairs lie within a Euclidean distance of 1 from each other, and only 9 pairs lie within a higher Euclidean distance from each other, attesting to the similarity in community composition between the pairs originating from the same sample.

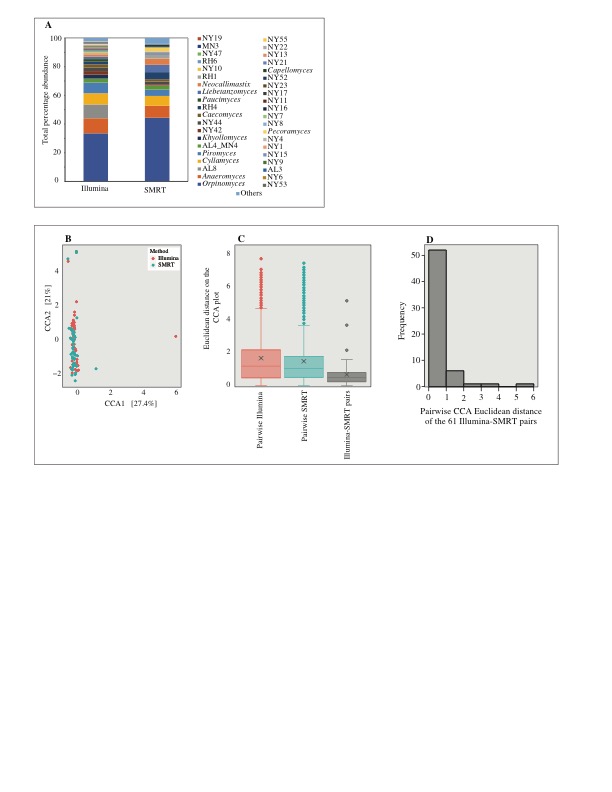

**Figure S5.** **Confirmation of the unique position of novel AGF genera (identified using D2 LSU amplicons and Illumina sequencing) using longer D1/D2 LSU amplicons and PacBio sequencing.** Maximum likelihood phylogenetic tree constructed using the alignment of the D1/D2 region from representatives of all cultured (blue) and uncultured (orange) genera, the D1/D2 region from representatives of the 49 novel genera (green) identified by PacBio sequencing in this study, and the D2 region from representatives of the 7 novel genera that were not identified in the PacBio dataset. Clades of genera are color coded by family as shown in the labels around the tree (for the newly proposed families *Neocallimastigaceae*, *Caecomycetaceae*, *Piromycetaceae*, and *Anaeromycetaceae*). Putative novel families encompassing multiple of the novel genera identified here, as well as genera previously-unaffiliated with the above four families are shown in red labels around the tree. These include novel family affiliated with the genus *Khoyollomyces*, novel family affiliated with the genus *Joblinomyces*, novel family affiliated with the genera *Buwchfawromyces* and *Tahromyces*, novel family affiliated with the genus *Aklioshbomyces*, and novel family affiliated with the genus *Paucimyces*. Bootstrap support is shown as black dots for nodes with >70% support.

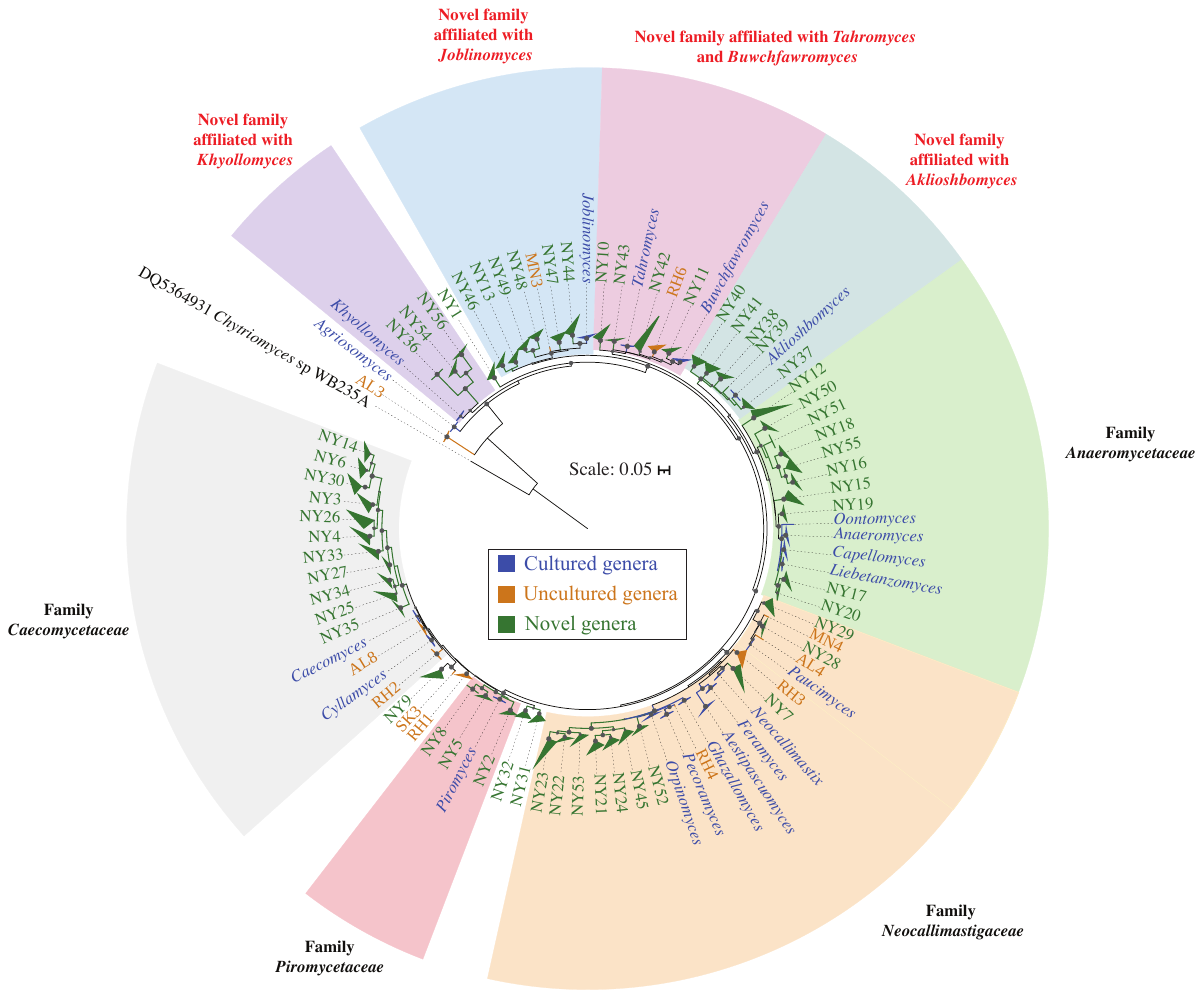

**Fig. S6. Patterns of AGF alpha diversity.** Samples with at least 1000 sequences were included (n=421); and the analysis was repeated with and without randomly subsampling 1000 sequences from each sample. Alpha diversity patterns were assessed using four different indices (observed number of genera, Shannon, Simpson, and Inverse Simpson) and the two sampling strategies. (A) Box and whisker plots showing the distribution of 4 alpha diversity measures for different animal species (top row), animal families (middle row), and animal gut types (bottom row). (B) Results of ANOVA showing the significant effect of the host species, animal family, animal gut type, but not domestication status, on alpha diversity measures regardless of the index used or the subsampling approach (p<0.0002). (C) Tukey test results for pairwise animal species, animal family, and animal gut type comparisons (8 comparisons each; 2 subsampling approaches x 4 diversity indices). Specifically, hindgut animals harbored a significantly less diverse community compared to foregut ruminants (in all 8 comparisons; p-value <0.00001). Accordingly, members of the hindgut family Equidae harbored a less diverse community when compared to the foregut ruminant families Cervidae and Bovidae (in 6/8 comparisons; p-value <0.002, and in 6/8 comparisons; p-value <0.00004, respectively); and horse communities were significantly less diverse than these of deer (6 /8 comparisons; p-value < 0.04), cows (in 6/8 comparisons; p-value < 0.00004), goats (6 out of 8 comparisons; p-value < 0.02), and sheep (4 out of 8 comparisons; p-value < 0.01). Within foregut ruminants, communities in animals belonging to the families Cervidae and Bovidae were not significantly different (7/8 comparisons; p-value >0.09). As well, on the animal host species level, most comparisons indicated no significant differences in diversity between deer, goat, cows, and sheep.

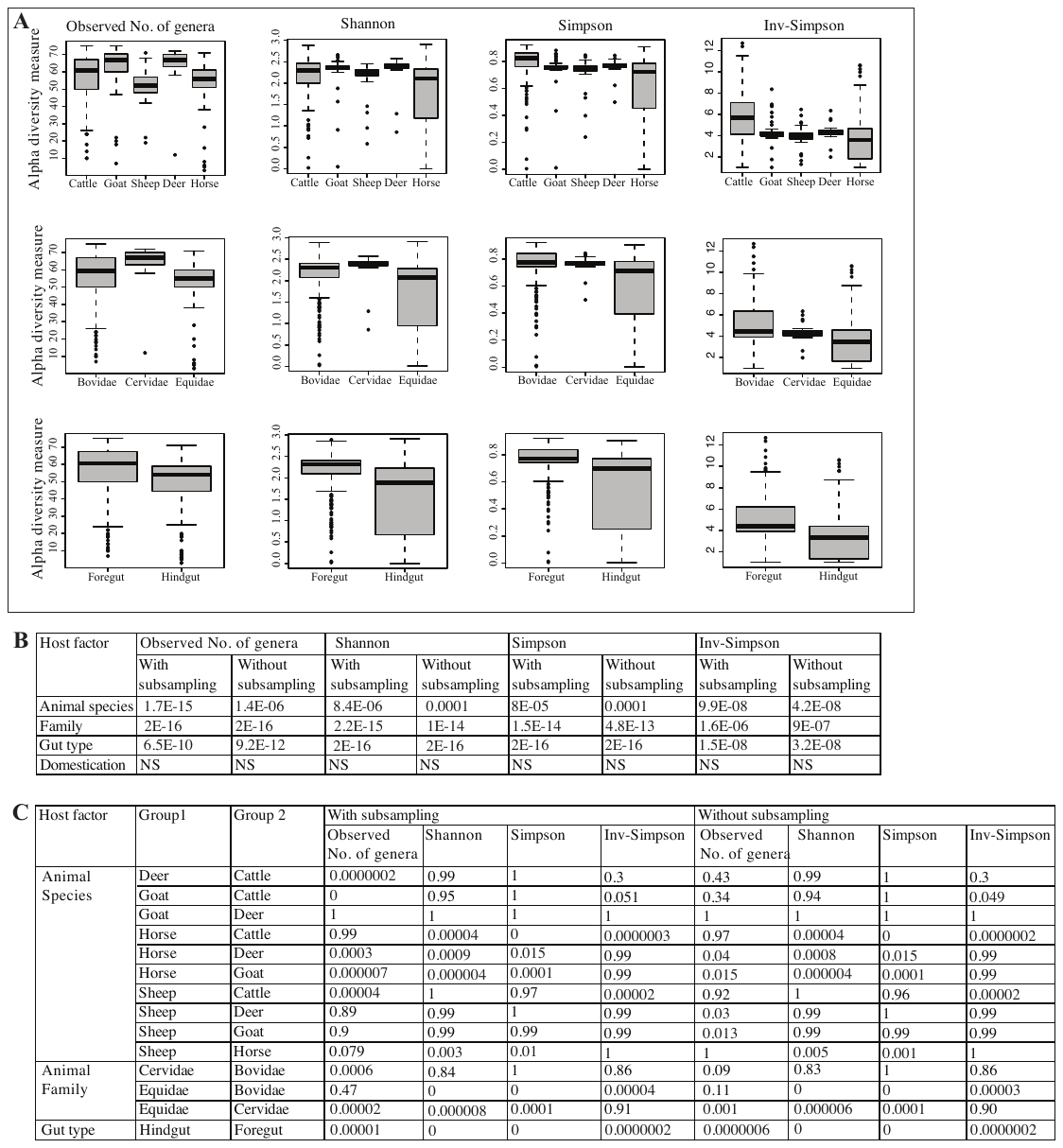

**Figure S7.** Box and whisker plots of the distribution of PCoA ordination distance of each sample to its group centroid. The PCoA plots are shown in Fig. 4a. Centroids were calculated using the command betadisper in the Vegan package for each animal species (left, only for animals with 4 or more individuals), animal family (middle), and animal gut type (right). Ordination distance between each point and the group centroid was calculated as the Euclidean distance between two points in an ordination plot. In general, samples clustered by the animal species (small variation in Euclidean distance to the animal species centroid), with only a few animals (e.g. horse, donkey, sheep, goat) showing large variation from their respective animal species centroid, usually due to only a few divergent samples. Interestingly, the microbial community structure showed a higher level of “variability” in hindgut animals when compared to foregut. Samples from animals belonging to the foregut families Bovidae, Cervidae, Giraffidae, and Camelidae clustered close to their respective animal family centroid. However, large variation was observed for the hindgut families Equidae, and Caviidae. This was also observed when comparing samples to their respective gut type centroid, where samples from foregut animals clustered close to their gut type centroid, while samples from hindgut animals showed large variation in their distance from the hindgut centroid. (B) Quantitative assessment of host factors affecting community structure via multivariate regression methods (multiple regression of matrices (MRM), Mantel tests for matrices correlations, and Procrustes rotation). Methods compared the AGF community dissimilarity matrix (Unifrac weighted (W), Unifrac unweighted (UW), Bray-Curtis, and Jaccard), to a matrix of each of the host factors tested (animal species, animal family, animal gut type, and domestication status). For the MRM analysis, results of the whole model (without partitioning variances into different host factors) are shown on top. The model was found to be significant regardless of the index used. For each of the three methods, the significance of correlation (depicted as the test p-value), and the degree the host factor is affecting community structure (depicted by the R^2^ regression coefficients of the MRM analysis, the Spearman correlation coefficients of the Mantel test, and the symmetric orthogonal Procrustes statistic of the Procrustes analysis) are shown. Significant p-values (<0.05) are shown in red text. Results of matrices correlation (12 total correlations; 3 methods x 4 dissimilarity indices) using each of the three methods, and regardless of the index used, confirmed the importance of animal host species, family, and gut type in explaining the AGF community structure. Animal host species and family were both found to be significant in all 12 correlations (p-value =0.001 for animal species, and <0.025 for animal family), while the animal gut type was found to be significant in 10 out of the 12 correlations (p-value <0.025). Further, comparing the correlation coefficients produced by each of the methods showed that the animal species explains more of the community structure (as evident by the higher R^2^ regression coefficients of the MRM analysis, the higher Spearman correlation coefficients of the Mantel test, and the higher symmetric orthogonal Procrustes statistic of the Procrustes analysis) than the animal family, or the gut type. This was true for 10 out of the 12 correlations. On the other hand, domestication status was only found significant in 3 out of the 12 total correlations, albeit with very low correlation coefficients.

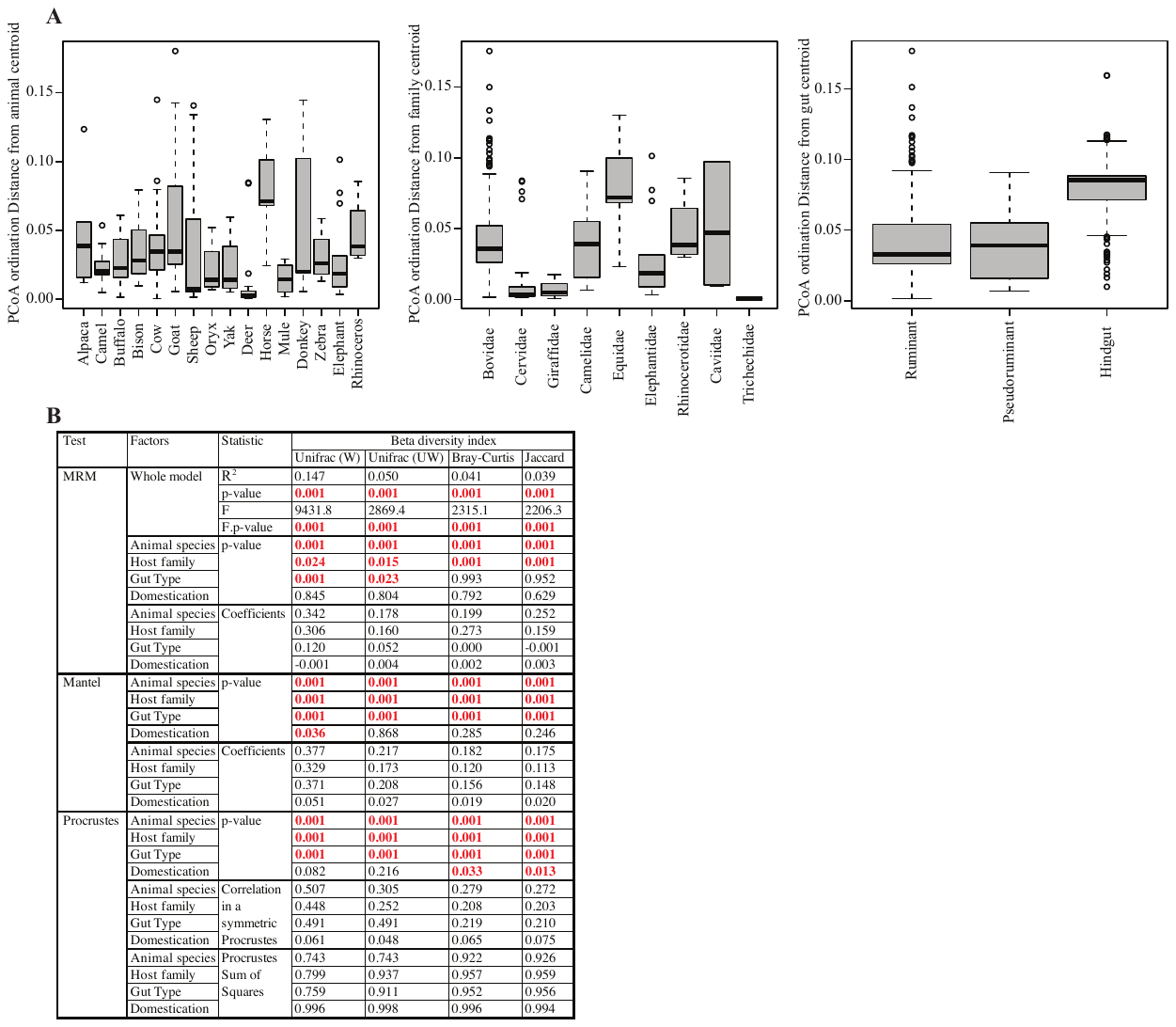

**Figure S8: Fungal genera-animal host preferences.** Double principal coordinate analysis (DPCoA) biplot constructed using the genera with abundance in the top 25% (n=28 genera). DPCoA uses both abundance and phylogenetic information about the samples, allowing both the samples and the taxa to be plotted on the same coordinate space, and thus the ordiantion distance between samples or their centroids and AGF genera could be compared. AGF genera with ordination distances close to sample centroids are abundant in these samples, and, therefore, contribute more to the community structure. For ease of visualization, individual samples are removed and only the animal species (hexagons, only for animals with 4 or more individuals), animal family (squares), and animal gut type (X) centroids are plotted, with standard deviation data ellipse shown for only the gut type. Centroids and data ellipses were generated using the command betadisper in the vegan package. Animals are color coded by their respective family as shown in the key and colors follow the same scheme as in Fig 1d. The AGF genera are shown as purple circles. The first two axes explained 69.4% of the variance. There was a clear separation of the hindgut families Equidae (orange square centroid), Rhinocerotidae (green square centroid), from the foregut families Bovidae (pink square centroid), Cervidae (yellow square centroid), and Giraffidae (red square centroid), with the pseudoruminant family Camelidae (purple square centroid) occupying an intermediate position. Of the 28 most abundant genera, 14 fell within the foregut ruminant ellipses, all of which were shared with the overlapping foregut pseudoruminant ellipse. Three additional genera fell within the foregut pseudoruminant ellipse. Only 9 genera fell within the hindgut ellipse outside the foregut ruminant and pseudoruminant ellipses. These genera included the Equidae-specific *Khoyollomyces* and AL3, the Trichechidae-specific *Paucimyces*, and the Rhinocerotidae-specific NY15. The genera *Orpinomyces*, and NY53 fell outside all ellipses, consistent with their high LIPA values of association with almost all families, hence their intermediary position on the biplot. Several genera, NY42, NY6, NY7, AL4/MN4, *Cyllamyces*, and AL8, fell within the ellipses of all gut types. These genera had moderate LIPA association values with almost all animal genera. The two genera *Piromyces*, and *Caecomyces* also fell within the ellipses of all gut types, but their position is most probably due to their strong association with the family Elephantidae.

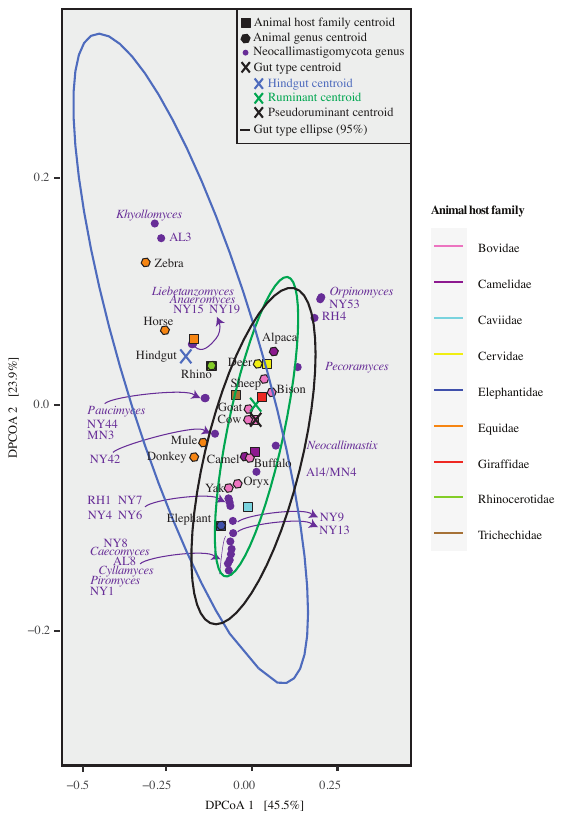

**Figure S9:** **Effect of biogeography, age, and sex on AGF community structure in cattle (A, E, I), goats (B, F, J), sheep (C, G, K), and horses (D, H, L).**

For testing the effect of biogeography, age, and sex on community structure, we opted to carry out comparisons only on samples belonging to the same animal species in an attempt to control for other host-associated factors that might conflate the results. For these comparisons, only the four most-sampled animal species (cattle, goats, sheep, and horses) were considered. We calculated within-animal Bray-Curtis dissimilarity indices for each animal subset and used them to construct PCoA plots (using the ordinate and plot_ordination commands in the Phyloseq R package) to describe the similarity between communities from the same animal genus originating from different geographical locations, or exhibiting different ages or sexes. The first two PCoA axes are plotted, and the percentage variance explained by each axis is shown for each plot. Samples are color-coded (as shown in the figure color key) by their country of origin (A-D), age (E-H), and sex (I-L). For biogeography effects, samples originated from different geographical locations (Argentina, Austria, Czech Republic, Egypt, Germany, Italy, Nepal, New Zealand, and the USA). For age, animals were classified as young (< 1 year), or adult (>1 year), and sex was either male or female. Samples where this information was not available are shown as NA. The first two axes explained 46.1%-55.5% of the variance depending on the animal subset. (M) To test for the significance of the above three factors in describing AGF community structure in each animal genus, PERMANOVA tests were run using the vegan command Adonis. The F-statistics p-value was used to assess the significance of AGF community difference between countries, young versus adult animals, and males versus females, and the sum of squares was used to assess the percentage variance explained by the country of origin for each of the four animal species. PERMANOVA tests showed that the country of origin explained 3.9% of variance in cattle AGF communities (F test p-value=0.002), 5.6% of variances in horses AGF communities (F test p-value=0.012), 23.62% of variances in goats AGF communities (F test p-value=0.001), and 30.84% of variances in sheep AGF communities (F test p-value=0.001). On the other hand, age explained 3.5-32.7% of variances, and sex explained 2.06-15.03% of variances in community structure.

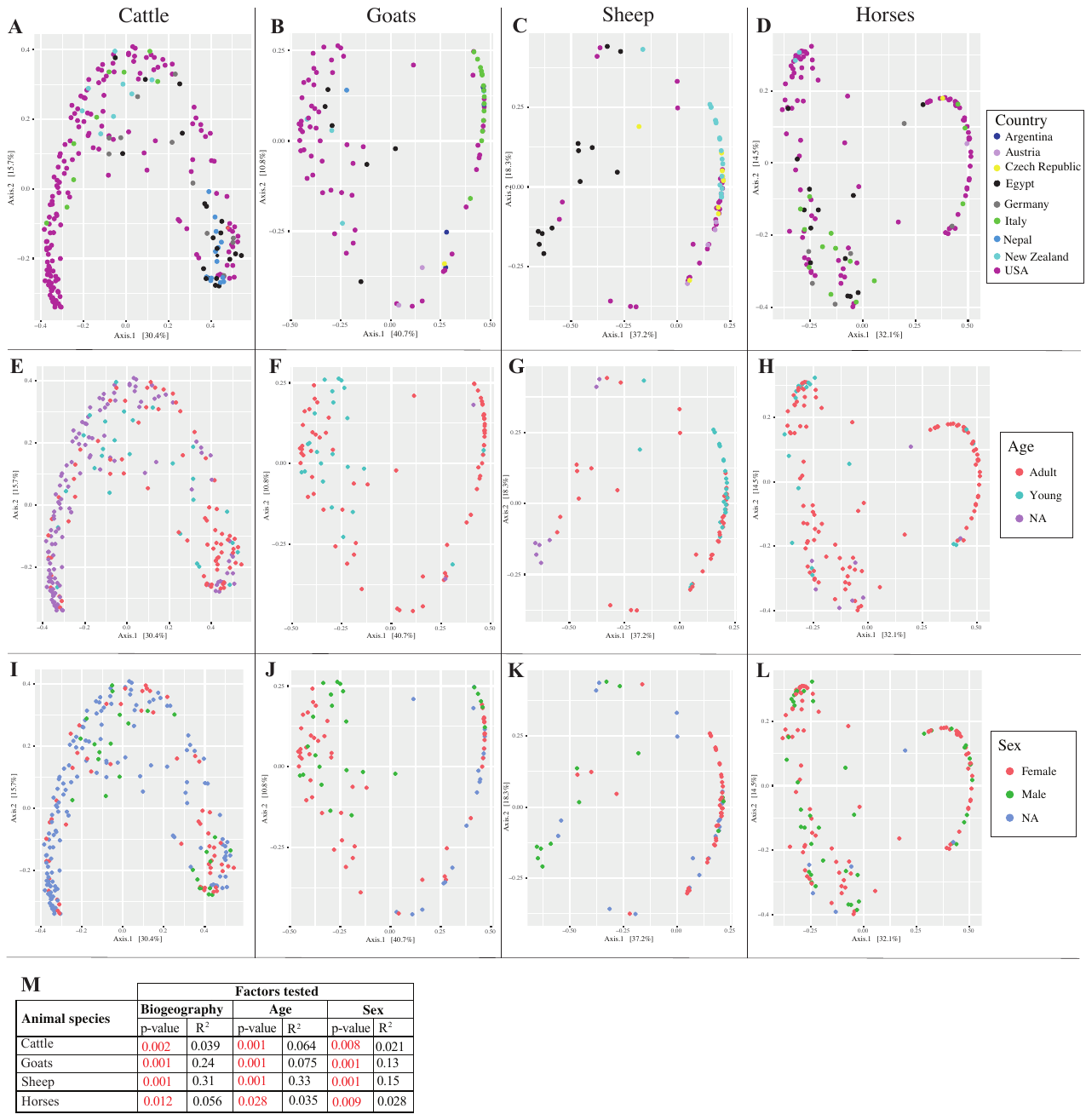

**Figure S10:** **Effect of amplicon length/region on pairwise sequence divergence estimates.** Comparison of all possible pairwise sequence identities of a group of 206 reference sequences, when the whole D1/D2 region was used (as would be obtained by SMRT sequencing using the NL1 forward primer/ GGNL4 reverse primer, X-axis) and when only the D2 region was used (as would be obtained by Illumina sequencing using primers employed here, Y-axis). Sequences were first aligned in Mafft, and the alignment was used to calculate pairwise distances in Mega. Aligned long sequences covering the D1/D2 regions were then trimmed in Mega to remove the D1 region, and this truncated alignment was then used to calculate pairwise distances in Mega.

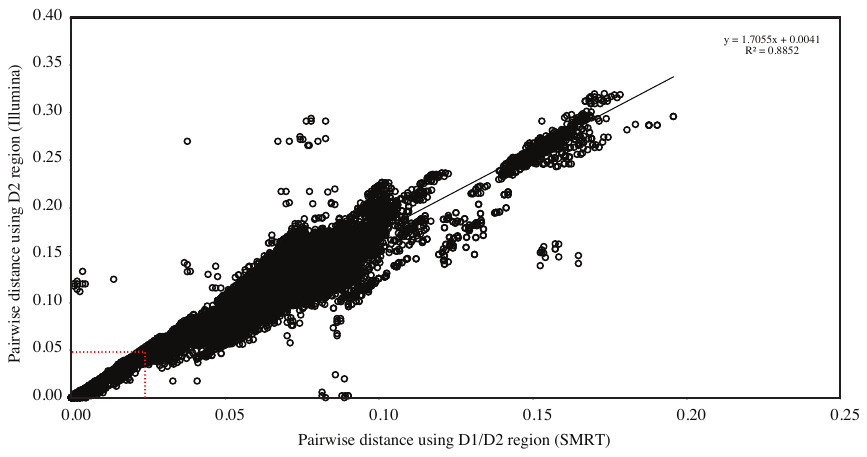

**Supplementary tables.**

**Table S1. Summary of previous high throughput culture-independent studies examining AGF diversity in herbivores.**

**Table S2: Metadata on all 661 datasets examined in this study**. Samples are grouped by their gut type, then animal host family, then animal host species. Country of origin (and state within USA), domestication status, and various metadata (including feed type, sex, and age) are also shown. Samples on which additional SMRT sequencing was conducted are highlighted in Red.

**Table S3.** **AGF genus-level community composition and Good’s coverage for the datasets studied.** Samples are shown in the same order as in Table S2. Samples on which additional SMRT sequencing was conducted are highlighted in Red.

**Table S4. Results of Wilcoxon test of significance for the distribution of the percentage of novel genera between different animal species (A), animal families (B), animal gut types (C), domestication status (D), and frequency of study (E).**

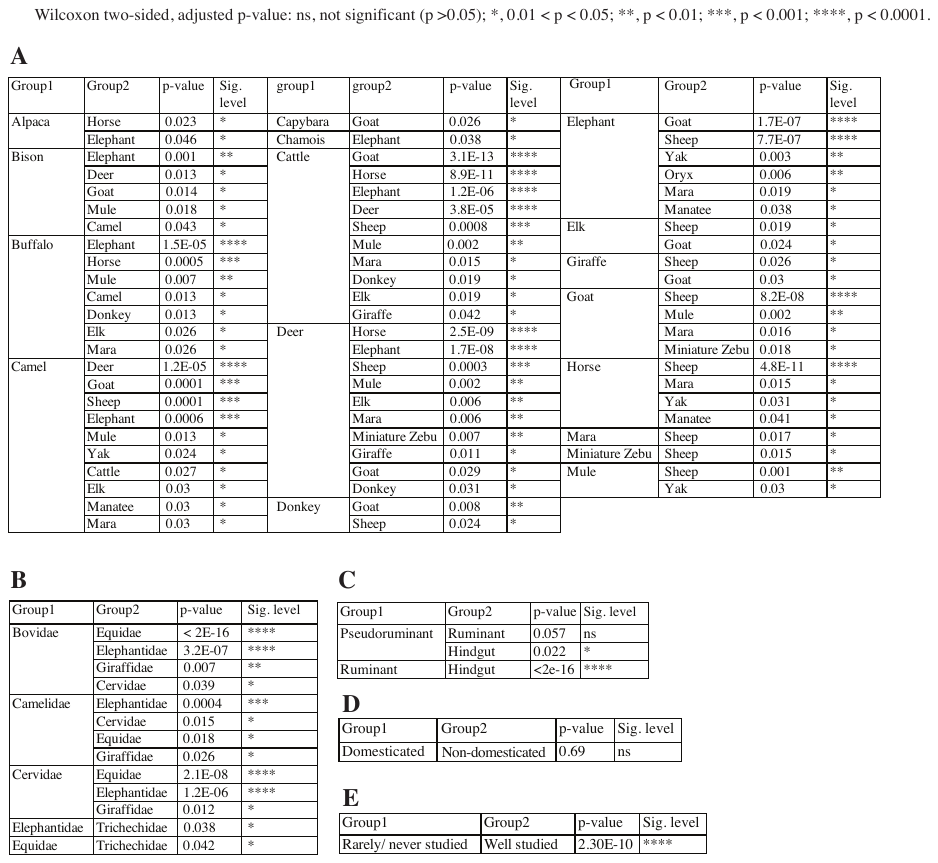

**Table S5. Distribution patterns of novel genera identified in this study*.**

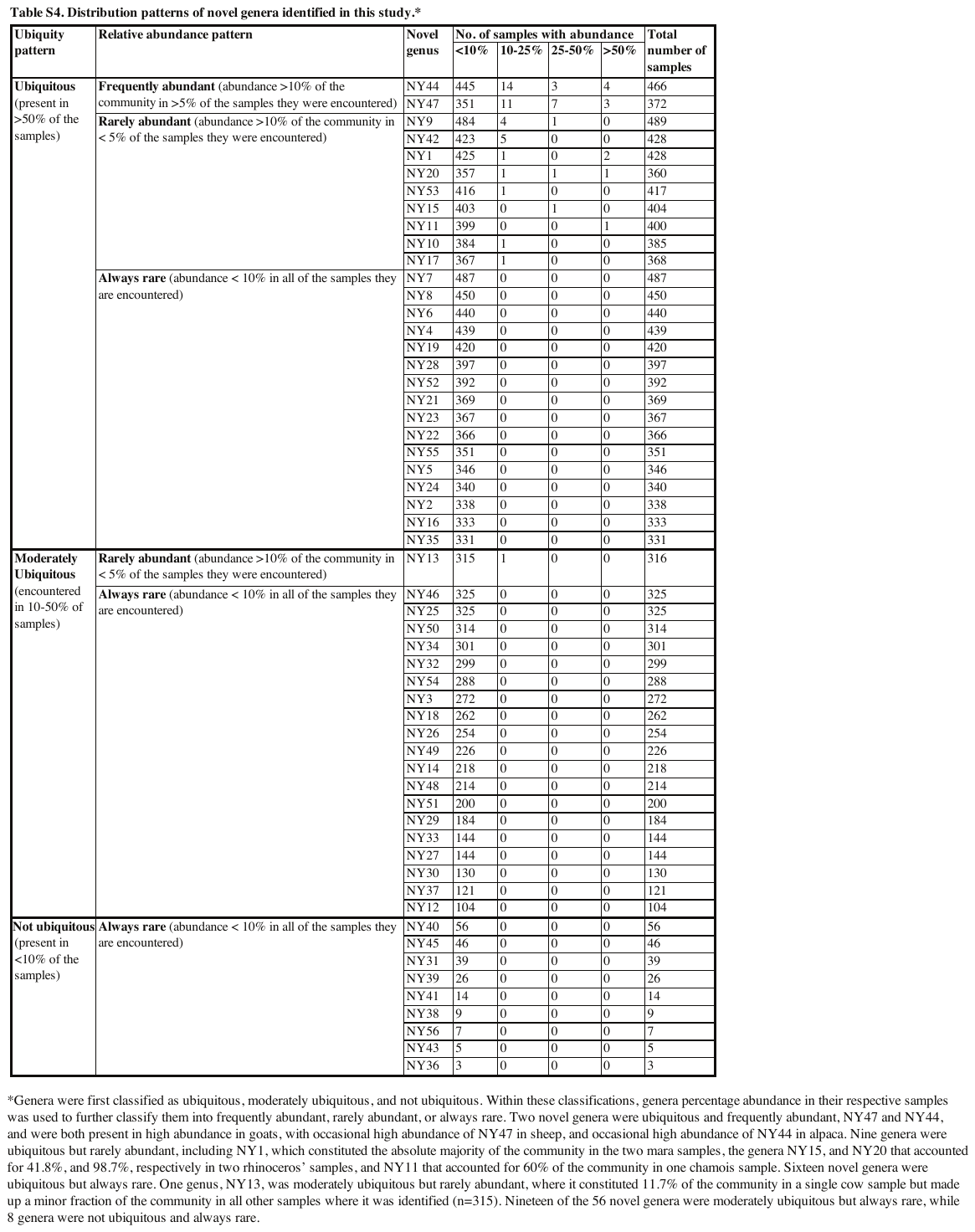

**Table S6. PacBio-generated sequences belonging to 49 of the 56 novel genera generated in this study, their novel genus affiliation, and their corresponding GenBank accession number.**

**Table S7.** **Topology comparison between trees constructed using Illumina-generated sequences affiliated with novel genera versus SMRT-generated sequences affiliated with novel genera.** The novel genera are color coded to reflect congruency between the two trees (yellow, positions congruent; green, positions non-congruent). Blue highlights the 7 novel genera that were missing from the PacBio dataset. These genera exhibited an extremely rare occurrence in the corresponding Illumina-sequenced samples (never exceeding 0.1% in any of the 61 samples), as well as the total dataset (abundances ranging between 0.0004-0.07% in the total Illumina dataset).

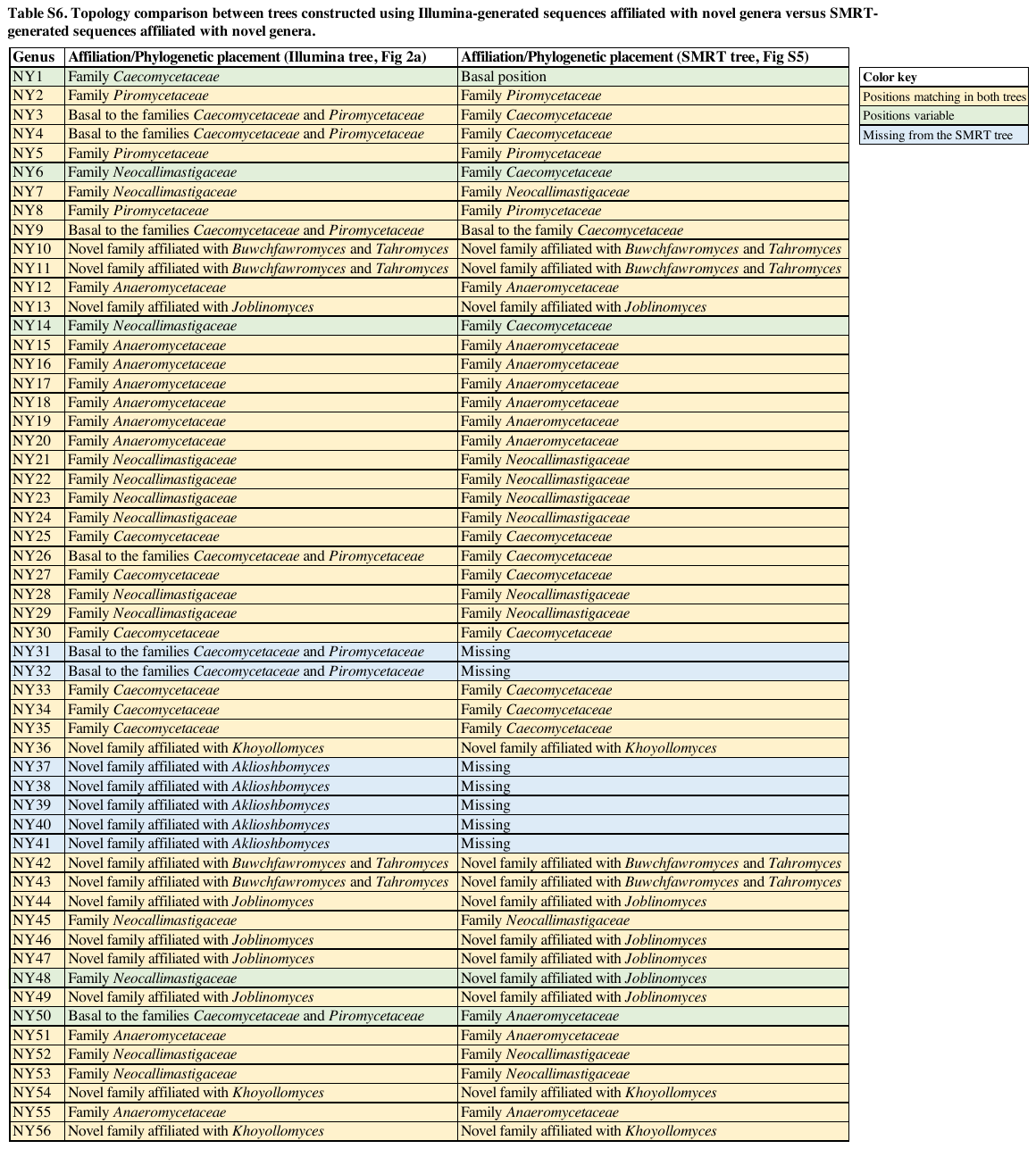

**Table S8.** Values of normalized stochasticity ratios (NST) calculated using the two indices Bray-Curtis and Jaccard.

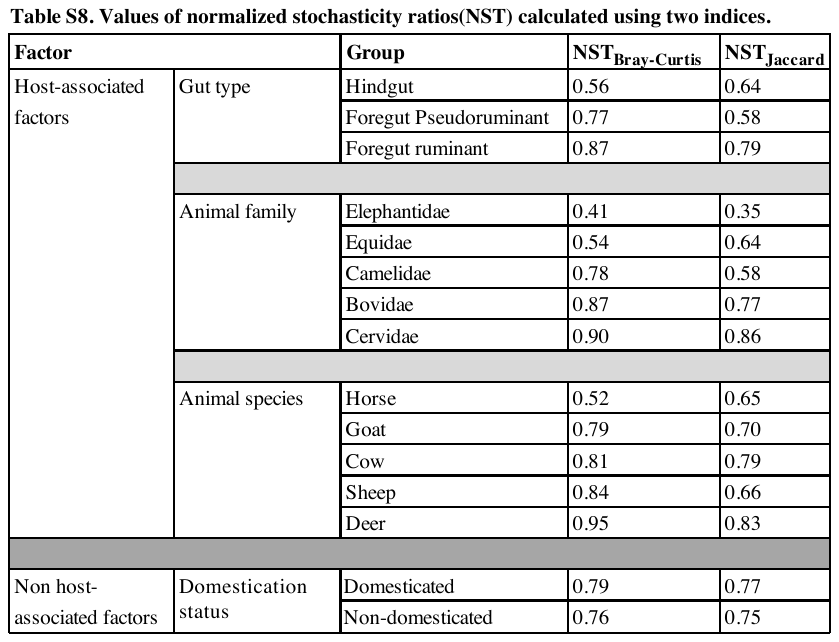

**Table S9. Results of Wilcoxon test of significance for the distribution of PACo residuals between different animal species (A), animal families (B), and animal gut types (C).**

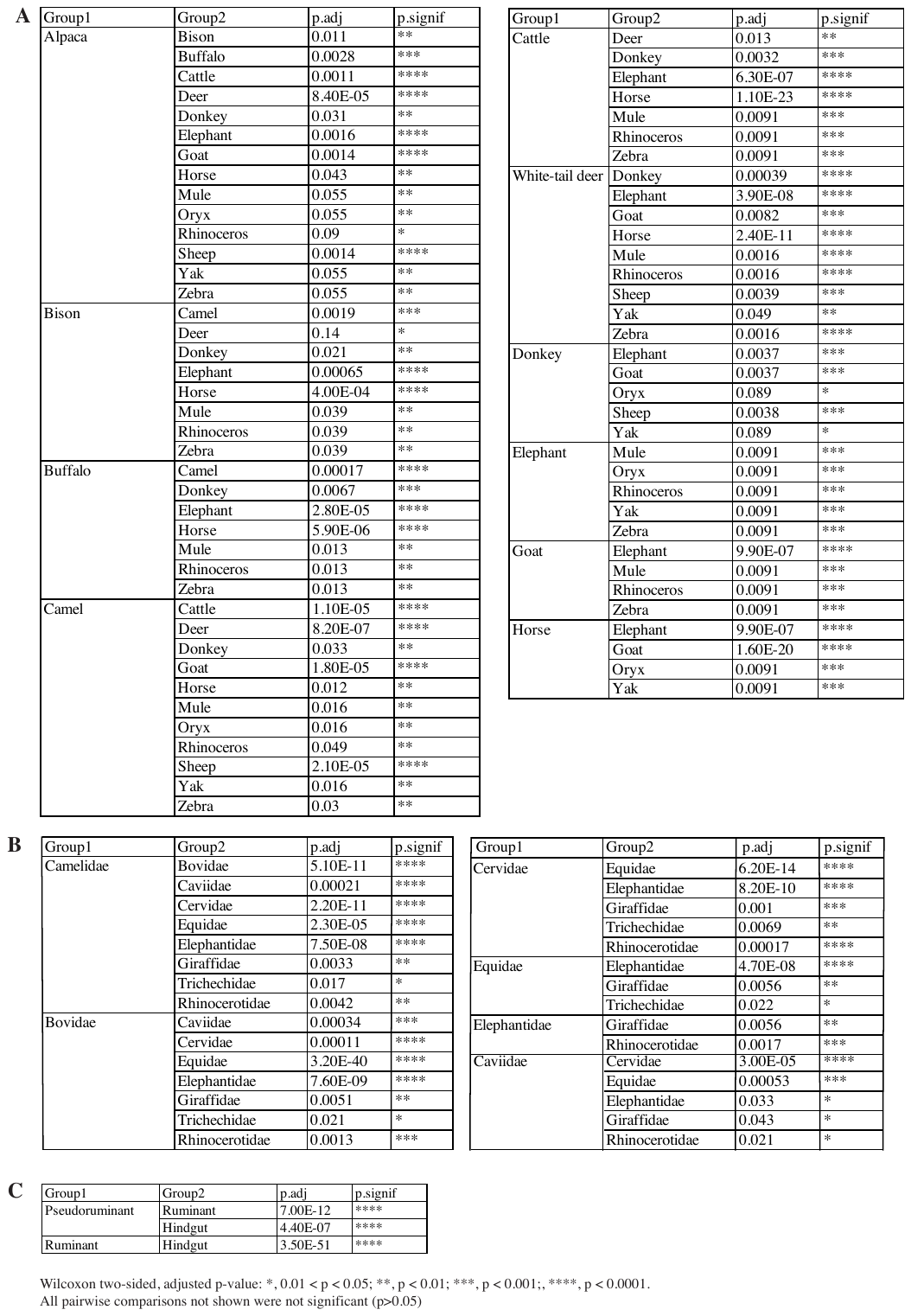

**Table S10.** Values of three global phylogenetic signal statistics and their associated p-values for the 37 AGF genera with significant correlations to the host phylogenetic tree*.

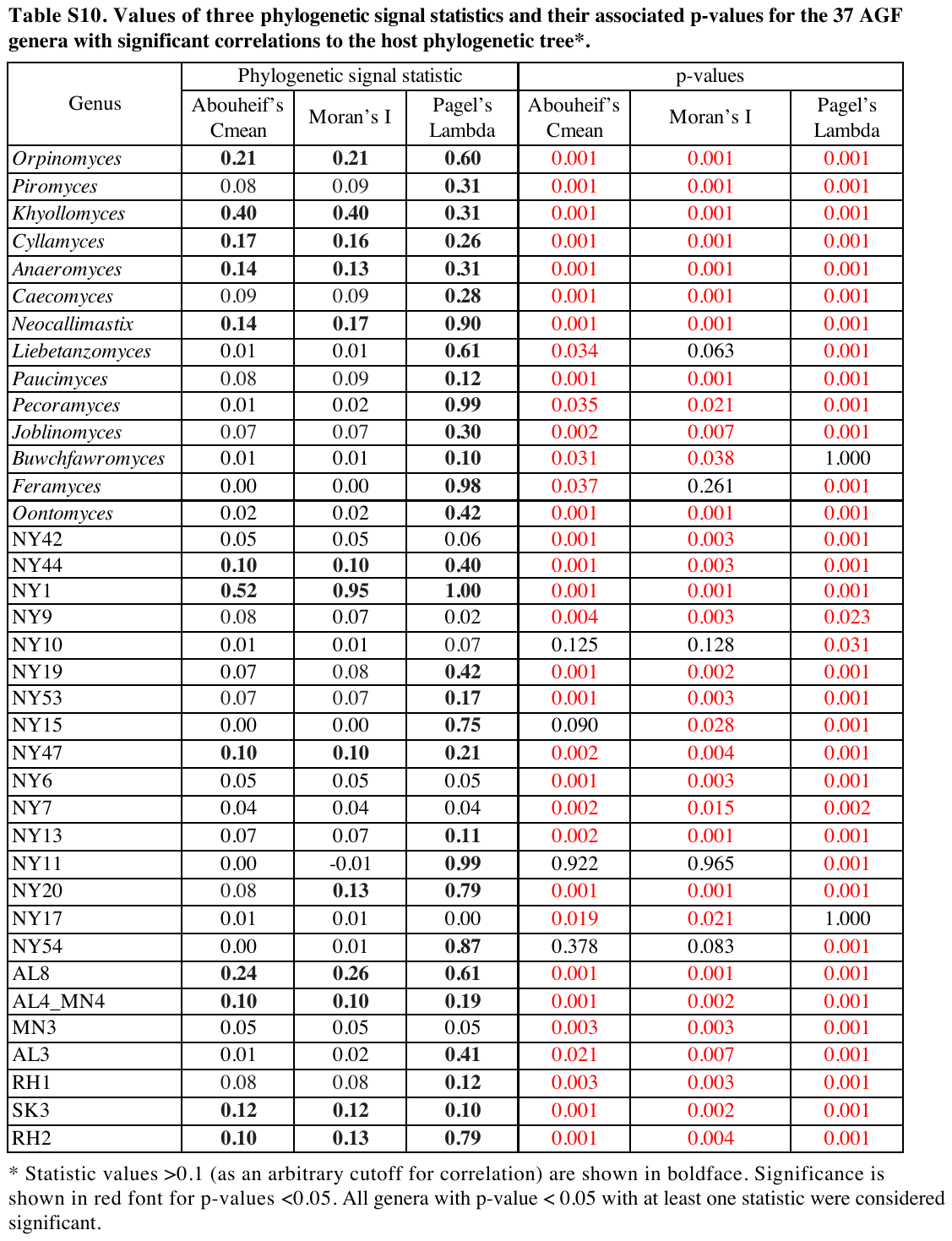

**Table S11.** Significant associations of AGF genera with studied animals based on LIPA values*. Note the high number of strong associations with hindgut animals, and the relatively lower number of strong host-AGF associations in ruminants (only in 3/22 animals: NY19 in bison, RH2 in oryx, AL8 in buffalo, NY9, SK3, and *Caecomyces* in yak, and *Neocallimastix* in elk). However, this lack of strong LIPA signal in foregut fermenters is countered by the identification of multiple intermediate and weak cophylogenetic signals (LIPA values 0.2-1; yellow in heatmap) per animal. For example, goats show intermediate and weak LIPA signals with *Joblinomyces*, NY47, and NY44, sheep show intermediate and weak LIPA signals with NY53 and *Orpinomyces*, buffaloes show intermediate and weak LIPA signals with *Cyllamyces*, NY9, NY14, and *Khoyollomyces*, bison show intermediate and weak LIPA signals with *Orpinomyces*, AL8, and RH1, oryx show intermediate LIPA signals with *Caecomyces*, *Buwchfawromyces*, and AL4/MN4, yak show weak LIPA signals with *Anaeromyces*, AL4, and *Orpinomyces*, deer show intermediate LIPA signals with *Orpinomyces*, NY13, and NY53, and elk show intermediate and weak LIPA signals with *Orpinomyces*, *Caecomyces*, *Khoyollomyces*, AL8, AL4, NY53, NY19, NY42, and NY7.

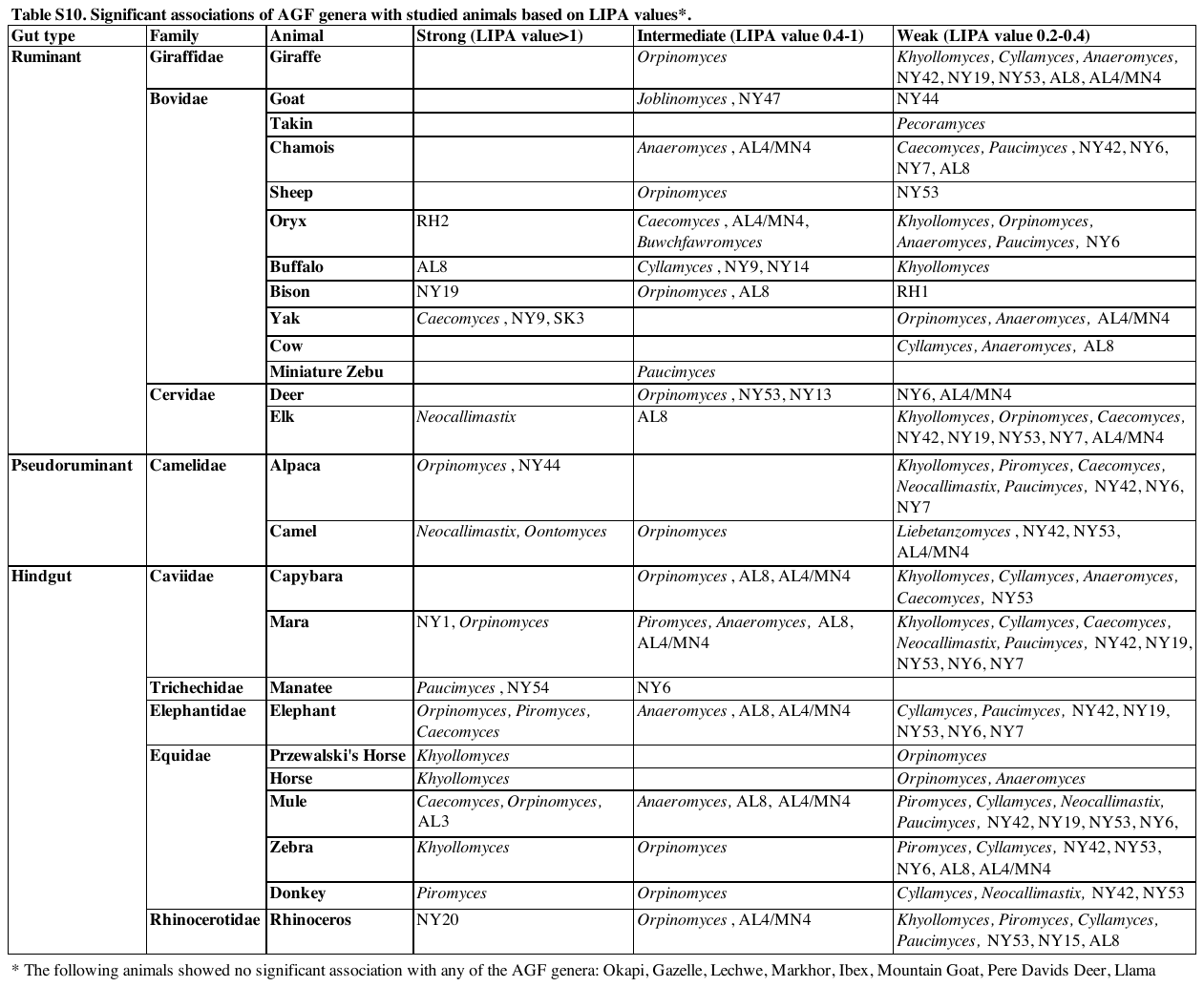

**Table S12.** **Effect of biogeography (country of origin), age, and gex on the AGF alpha diversity in cattle, horses, goats, and sheep.** (A) ANOVA results for the effect of biogeography (country of origin), age, and sex on alpha diversity measures (Shannon, observed number of genera, Simpson, and Inverse Simpson diversity indices). Significant p-values (p<0.05) are shown in red text. ANOVA showed that the country of origin had a significant effect on alpha diversity in cattle (with all indices, p-value <0.03), horses (with 3 out of 4 indices, p-value <0.04), but not in goats (with 3 out of 4 indices, p-value >0.1), or sheep (with 3 out of 4 indices, p-value >0.05). On the other hand, animal sex largely had no significant effect on alpha diversity (p-value>0.05). Animal age only showed significant effect on the alpha diversity of horses (with all indices, p-value <0.03), goats (with all indices, p-value <0.01), and sheep (with 2 out of 4 indices, p-value <0.003), but not cattle. Since out of the three non-host associated factors tested, biogeography showed the most significant effect on alpha diversity, we further carried out Tukey HSD tests for pairwise country (B), and US state (C) comparisons for the two animals with the most significant results (cattle and horses). Cattle samples from the USA, and New Zealand were found to be more diverse than cattle from Germany, but no significant difference was observed in alpha diversity of cattle from all other countries. Similarly, horse samples from USA were found to be more diverse than horses from Germany. Within the USA, the state of origin significantly affected the alpha diversity in cattle, and horses. Analysis of variance showed only significantly higher diversity in cattle originating from OK, in comparison to AZ, CT, and FL, and significantly higher diversity in horses originating from OK, in comparison to CT.

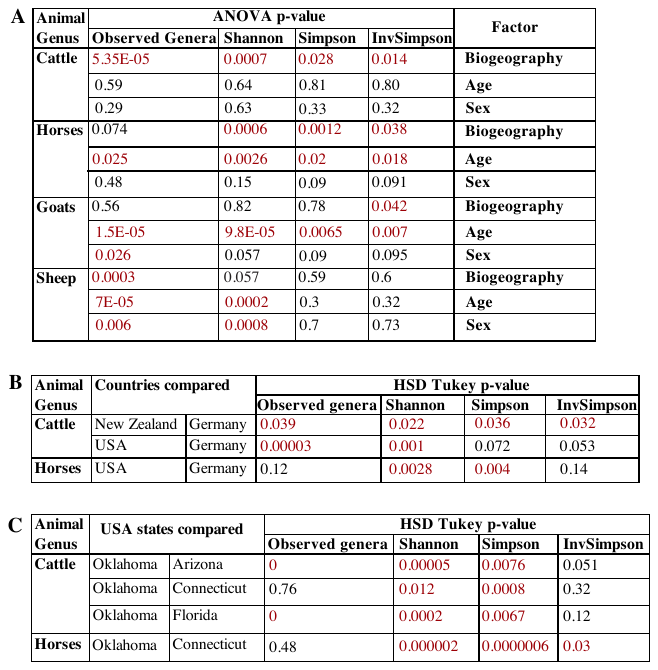
